# How much warming can mosquito vectors tolerate?

**DOI:** 10.1101/2024.01.03.574109

**Authors:** Lisa I. Couper, Desire Uwera Nalukwago, Kelsey P. Lyberger, Johannah E. Farner, Erin A. Mordecai

**Affiliations:** UC Berkeley; Stanford University

## Abstract

Climate warming is expected to substantially impact the global landscape of mosquito-borne disease, but these impacts will vary across disease systems and regions. Understanding which diseases, and where within their distributions, these impacts are most likely to occur is critical for preparing public health interventions. While research has centered on potential warming-driven expansions in vector transmission, less is known about the potential for vectors to experience warming-driven stress or even local extirpations. In conservation biology, species risk from climate warming is often quantified through vulnerability indices such as thermal safety margins – the difference between an organism’s upper thermal limit and its habitat temperature. Here, we estimated thermal safety margins for 8 mosquito species (including *Aedes aegypti* and *Anopheles gambiae*) that are the vectors of malaria, dengue, chikungunya, Zika, West Nile and other major arboviruses, across their known ranges to investigate which mosquitoes and regions are most and least vulnerable to climate warming. We find that several of the most globally important mosquito vector species have positive thermal safety margins across the majority of their ranges when realistic assumptions of mosquito behavioral thermoregulation are incorporated. On average, the lowest climate vulnerability, in terms of both the magnitude and duration of thermal safety, was just south of the equator and at northern temperate range edges, and the highest climate vulnerability was in the subtropics. Mosquitoes living in regions largely comprised of desert and xeric shrubland biomes, including the Middle East, the western Sahara, and southeastern Australia have the highest climate vulnerability across vector species.

## Introduction

Climate warming is already causing shifts in the distribution and species interactions of many taxa, affecting ecosystem processes and health, and these effects are projected to accelerate with continued carbon emissions (Elsen *et al*. 2022; Grimm *et al*. 2013; Urban 2015). In particular, climate warming affects many ectotherms due to their temperature-sensitive life history and population dynamics, altering their geographic ranges and imposing thermal stress near their upper thermal limits (Deutsch *et al*. 2008; Parmesan 2006). Mosquitoes, which transmit many of the most important and high-burden infectious diseases worldwide, including malaria, dengue, Zika, and West Nile, are projected to experience changes in population dynamics, range limits, and transmission potential due to climate change (Harrigan *et al*. 2014; Kraemer *et al*. 2019; Ryan *et al*. 2019, 2021). Near cool range edges, warming is widely expected to promote mosquito population growth and disease transmission (Chen *et al*. 2010; Medlock & Leach 2015; Mordecai *et al*. 2019; Parmesan & Yohe 2003; Ryan *et al*. 2019). However, the impacts of warming across other portions of mosquito ranges are less clear and will depend on the extent of warming (*e*.*g*., the magnitude and duration of high temperatures), as well as species physiological limits and response strategies. In particular, ectotherm physiology theory and empirical work predict that upper thermal limits may restrict species persistence in currently suitable ranges under excessive warming (Deutsch *et al*. 2008; Huey & Berrigan 2001; Kearney *et al*. 2009a; Kellermann *et al*. 2012; Pinsky *et al*. 2019; Pörtner *et al*. 2006; Sunday *et al*. 2014). Understanding which mosquito species and regions will be most impacted by warming is critical for preparing targeted vector control and disease prevention strategies.

The potential impacts of climate change across species and broad spatial scales are often quantified through vulnerability indices such as ‘thermal safety margins’—the difference between an organism’s critical thermal maximum and its habitat temperature—(sometimes used interchangeably with ‘warming tolerance’; Angilletta 2009; Clusella-Trullas *et al*. 2021; Deutsch *et al*. 2008). Thermal safety margins, which reflect the amount of additional warming an organism could experience before trait performance is inhibited, have been estimated for hundreds of ectotherm species including lizards, snakes, fish, and insects (Deutsch *et al*. 2008; Diamond *et al*. 2011; Sunday *et al*. 2019, 2014; Vinagre *et al*. 2019). These prior studies have often found the lowest thermal safety margins, and thus the highest vulnerability to warming, for species in the tropics. From a conservation standpoint, this finding has raised concern given the high biodiversity contained in the tropics. However, whether this pattern of maximum thermal stress in the tropics applies for species that threaten human health, such as mosquitoes that vector pathogens, has not been rigorously evaluated.

Thermal safety margins can provide biased estimates if the estimated critical thermal maximum and/or environmental temperatures do not reflect those relevant to the organism (Clusella-Trullas *et al*. 2021). For example, air temperature in full sun, as is often captured by weather station data sources, can differ drastically from an organism’s body temperatures (Kearney *et al*. 2009b; Sunday *et al*. 2014). Similarly, environmental temperature estimates with coarse temporal resolution (*e*.*g*., daily or monthly scales) may fail to capture short-term thermal extremes, which can have major impacts on organismal fitness and population growth (Buckley & Huey 2016a, b; Dowd *et al*. 2015; Vasseur *et al*. 2014). To address these limitations, recent advances in climate monitoring and microclimate and biophysical modeling have enabled increasingly accurate estimates of realized environmental temperatures on time scales relevant to short ectotherm life cycles (Hersbach *et al*. 2020; Kearney & Porter 2017, 2020).

To date, no work has systematically estimated thermal safety margins for the mosquitoes that transmit major human pathogens. Here, we incorporate these recent advances in physiological and bioclimatic modeling to estimate thermal safety margins for 8 major mosquito vector species across their ranges to identify the disease systems and locations that are most and least vulnerable to climate warming. We specifically ask: how much buffering do mosquitoes have from thermal extremes in their current ranges? We define thermal safety margins as the difference between a species’ critical thermal maximum, estimated in the laboratory for a range of life history traits under constant temperatures, and the hottest hourly body temperature experienced in the coolest microhabitat available (Clusella-Trullas *et al*. 2021; Pinsky *et al*. 2019). To ensure that our estimates provide a meaningful metric of climate vulnerability, we explicitly: 1) incorporate the realistic potential for mosquito movement to cooler, fully-shaded microhabitats (*i*.*e*., ‘behavioral thermoregulation’); 2) estimate operative mosquito body temperature rather than air temperature, as these can differ markedly (Kearney *et al*. 2009b; Sunday *et al*. 2014; Woods *et al*. 2015); and 3) use hourly temperature data to capture the impact of short-term thermal extremes that may be missed with more temporally coarse temperature estimates (*e*.*g*., daily, monthly). Further, as mosquito climate vulnerability will depend not only on the difference in critical thermal maxima and body temperatures, but also how often and for how long critical thermal maxima are exceeded, we also estimated the time spent in thermal danger (here, the number of consecutive hours or days when body temperatures exceed critical thermal maxima).

## Methods

### Mosquito thermal tolerance data

We included the following 8 mosquito species in our investigation as these constituted major disease vectors with available thermal tolerance estimates: *Aedes aegypti, Ae. albopictus, Anopheles gambiae, An. stephensi, Culex annulirostris, Cx. pipiens, Cx. quinquefasciatus, and Cx. tarsalis* (Table 1). Together, this includes the major vectors of 15 of the most important human vector-borne pathogens (Table 1). As our measure of the estimated critical thermal maximum (CT_max_) for each species, we used the estimated thermal maxima (T_max_) calculated from thermal performance curves derived from laboratory experimental data on adult survival and synthesized in Mordecai *et al*. (2019), Villena *et al*. (2022), and Cruz-Loya *et al*. (2024) (Table 1). While for many species, thermal performance estimates are available for multiple life history traits (*e*.*g*., immature development rate, immature survival, biting rate), we used estimates for adult survival for all species for consistency and because it is likely the life history trait that is the most accurately and precisely estimated (*i*.*e*., because the decline in trait performance above the thermal optimum can be readily observed experimentally and estimated mathematically).

**Table 1.**
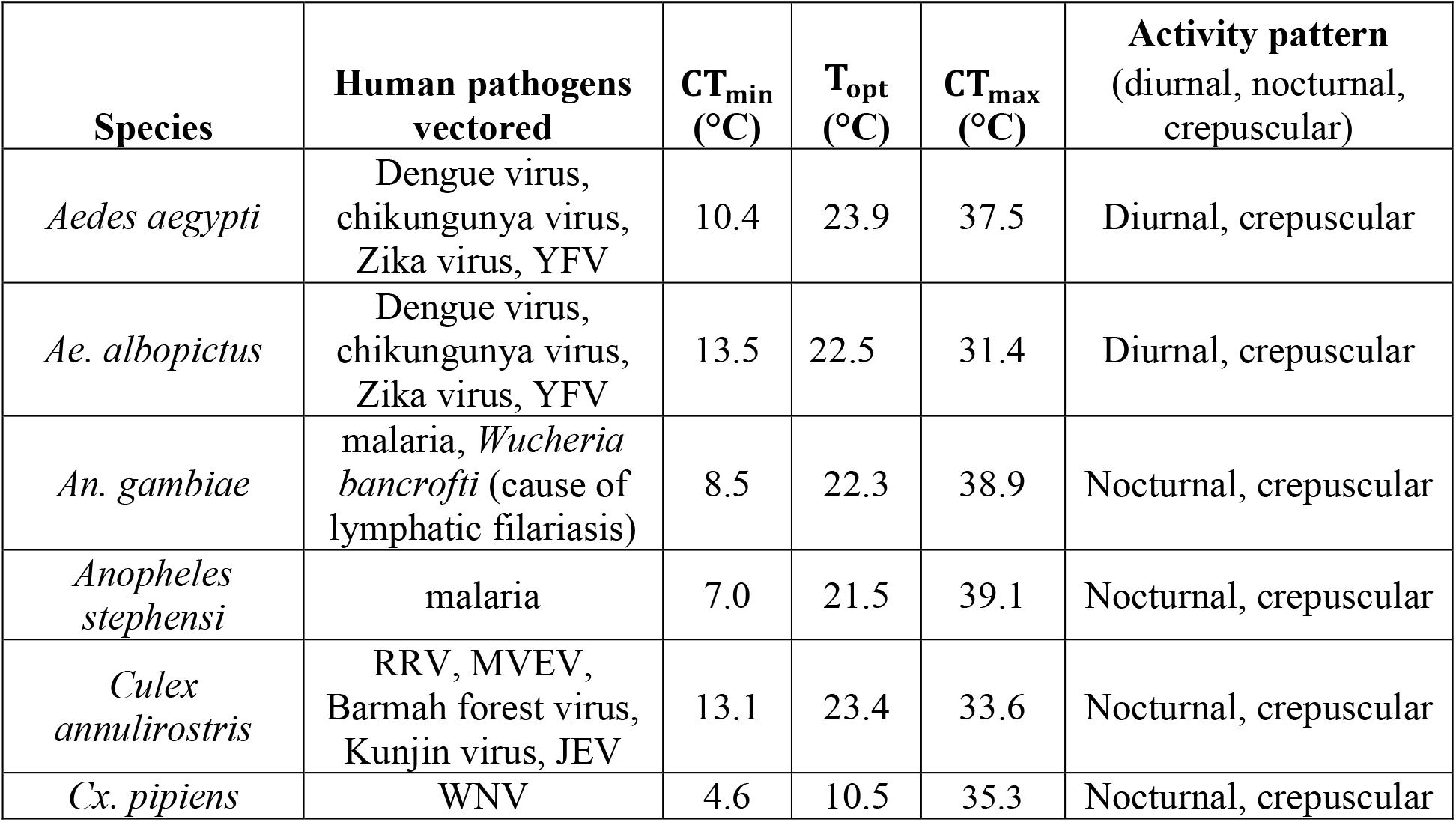
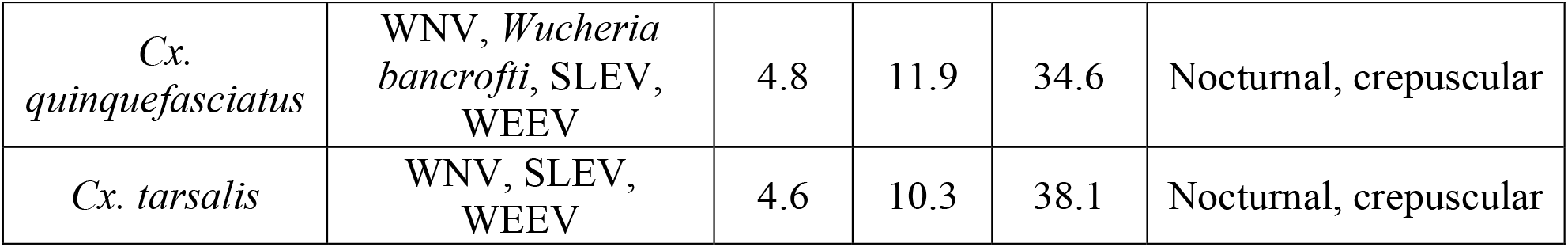
Species-specific trait values used in biophysical models of mosquito body temperature. For each species, the thermal limit values (CT_max_, CT_min_, and T_opt_) are for adult survival. The following abbreviations are used below: MDR for mosquito immature development rate, YFV for yellow fever virus, RRV for Ross river virus, WNV for West Nile fever, MVEV for Murray Valley encephalitis virus, JEV for Japanese encephalitis virus, SLEV for St. Louis encephalitis virus, and WEEV for Western equine encephalitis virus.

Further, it is likely the trait most strongly linked to overall fitness as mortality instantly and irreversibly reduces fitness to zero. Lastly, the immature life stages occur in water, where temperatures may differ substantially from that of the surrounding air, for which our environmental temperature estimates apply. These estimates of the critical thermal maximum (CT_max_), as well as the critical thermal minima (CT_min_) and optima (T_opt_) were derived from Bayesian thermal response models fit to empirical data on mosquito adult survival at different constant temperatures. As a sensitivity analysis, we estimated thermal safety margins for these 8 species using the critical thermal maximum from the most thermally tolerant life history trait (Supplemental Table S1, Supplemental Figure S1).

There is no consensus methodology for estimating organismal thermal tolerance, but rather a wide range of methods including static and dynamic heat tolerance assays (*e*.*g*., ‘thermal knockdowns’) of varying durations (Bates & Morley 2020; Clusella-Trullas *et al*. 2021; Hoffmann *et al*. 2013; Jørgensen *et al*. 2019; Lutterschmidt & Hutchison 1997; Rezende *et al*. 2014; Terblanche *et al*. 2007). Experimental methodology can impact the estimated CT_max_ (*e*.*g*., Terblanche *et al*. 2007; Woods *et al*. 2018), with longer duration heat stress assays—as in the constant temperature lab exposures from which our CT_max_ estimates are derived—typically yielding lower estimated CT_max_ than short duration assays (Bates & Morley 2020; Peck *et al*. 2009; Woods *et al*. 2018). However, thermal knockdown assays measure a single trait—the ability for a mosquito to stay standing and right itself at a given heat exposure—with an unknown linkage to key demographic, life history, and transmission-related traits; and these assays often measure time duration rather than temperature as their output. Moreover, these assays have not been conducted systematically across mosquito species. We instead chose to use CT_max_ values from constant temperature experiments because these capture performance that is meaningfully linked to mosquito population dynamics and distributions, but which may be lower than values derived from short-term knockdown experiments. For a given species, these empirical estimates often come from a lab colony representing a single, potentially lab-adapted population. Thus, although mosquito thermal tolerance can vary between populations, we applied the available thermal tolerance estimate to the species across its range.

### Mosquito occurrence data

To determine the distribution over which to estimate thermal vulnerability, we used published and/or publicly available occurrence records (*i*.*e*., a collection with an accompanying latitude and longitude) from the Global Biodiversity Information Facility (GBIF; see Supplemental Table S2 for full GBIF doi) for each species (Supplemental Figure S2). For species with hundreds of occurrence records (*i*.*e*., *Ae. aegypti*), we used a spatially stratified sampling approach to select 20-30 records from each 10° latitudinal band on which to estimate thermal safety margins. That is, within each 10° latitudinal band, we grouped records by longitude and randomly sampled from within each group to minimize overrepresentation in specific regions (*e*.*g*., cities or highly visited natural areas). While our approach was not designed to comprehensively cover the entire range of a given species, in cases of clear data gaps in GBIF (*e*.*g*., *Anopheles stephensi* in Kenya), we further supplemented these occurrence records with those from published data sources through targeted literature searches. We used the associated latitude and longitude to classify occurrence records into one of 14 biomes (Olson *et al*. 2001) and as ‘tropical’ (0-23.5°), ‘subtropical’ (23.5-35.0°), and ‘temperate’ (35.0-66.5°).

### Estimating mosquito body temperatures

For each vector species and collection location, we estimated mosquito body temperatures using NicheMapR: a suite of R programs for microclimate and biophysical modeling (Kearney & Porter 2020). In the first step, we used the microclimate model with hourly ERA5 weather data as input, to estimate microclimate conditions in both full shade and full sun (Kearney & Porter 2017). Specifically, the microclimate model input includes ERA5 data on wind speed, longwave radiation flux, solar radiation, surface pressure, 2 m temperature and 2 m dewpoint temperature, total precipitation, and cloud cover (Klinges *et al*. 2022). These data are available at a spatial resolution of 0.25° and are downscaled to an approximately 30 m resolution based on local terrain effects (*i*.*e*., slope, aspect, elevation). The downscaled microclimate conditions are then used as the input in the biophysical model, which is used to estimate steady-state body temperatures. In addition to the microclimate conditions, the biophysical model utilizes information on organismal functional traits including body size, daily activity patterns (*e*.*g*., diurnal, nocturnal, crepuscular), and thermal limits (Kearney & Porter 2020). We supplied species-specific parameter values for daily activity patterns and thermal limit estimates (*e*.*g*., point estimates and 95% credible intervals for T_opt_, CT_max_, CT_min_ for adult survival; Table 1, Supplemental Table S1). In the biophysical model, T_opt_ is considered the organisms’ preferred body temperature (*i*.*e*., the temperature that it will attempt to maintain), while CT_max_ and CT_min_ impact the organisms’ thermoregulatory behaviors (Kearney & Porter 2020). We assumed that all mosquito species had a body size of approximately 3 mg with 85% solar absorptivity (Brust 1967). Using these parameter values, we estimated species’ thermal safety margins with behavioral thermoregulation (*i*.*e*., the ability to move across fully exposed to fully shaded microhabitats; main results). Results without behavioral thermoregulation are presented in Supplemental Figure S3. These models have been extensively validated in a wide range of field conditions (Kearney *et al*. 2014; Kearney & Maino 2018; Kearney & Porter 2017).

### Incorporating impacts of drought on vector life cycles

In environments with highly seasonal precipitation, mosquito populations may reduce activity or aestivate during drought (Adamou *et al*. 2011; Lehmann *et al*. 2010), and thus avoid exposure to high temperature extremes experienced during that time. To incorporate potential seasonal and aestivation responses driven by low moisture availability, we masked out any periods in which the prior 30 or more days had soil moisture below 5%. That is, if the prior 30+ days each had soil surface wetness <5%, then hourly thermal conditions from that point onward were excluded from our thermal vulnerability calculations until soil moisture rose above 5% again. This soil moisture metric roughly corresponded to precipitation events (Supplemental Figure S4). Although some mosquito species may persist during drought by using artificial water sources (*e*.*g*., *Aedes aegypti*; Trpiš 1972), we used this drought mask for all species for consistency, as comprehensive information on the drought responses of mosquito populations across their ranges is not available. For all species, incorporating this drought mask had minimal impact on the thermal safety estimates (Supplemental Tables S4-S6, Supplemental Figure S5).

### Estimating thermal safety margins

To quantify mosquito vulnerability to climate warming, we calculated thermal safety margins: the difference between an organism’s critical thermal maximum (CT_max_) and the warmest temperature the organism experiences (T_e_) in the coolest microhabitat available (Deutsch *et al*. 2008; also referred to as ‘warming tolerance’). We interpret regions and species with low thermal safety margins as having higher thermal vulnerability, but do not attempt to draw specific cut-offs in what constitutes vulnerability based on this metric. By using highly resolved microclimate data (*i*.*e*., spatial resolution of approximately 30 m), and estimating T_e_ both with and without behavioral thermoregulation, we believe the thermal safety margins calculated here provide a relevant metric of thermal vulnerability. To capture both the magnitude and duration of thermal safety, we specifically calculated three related indices: 1) thermal safety margins, 2) the longest continuous period (in hours) that T_e_ exceeds CT_max_ (*i*.*e*., ‘thermal danger’), and 3) the longest streak of consecutive days in thermal danger (where a day is counted if T_e_ > CT_max_ for at least one hour). Together, these capture both average thermal safety and seasonal and diurnal variation in thermal safety. After estimating these indices at each collection location for each species, we used generalized additive models (GAM) to estimate latitudinal patterns of thermal vulnerability (Supplemental Methods). We fit all models using a smoothed function of up to eight knots estimated using restricted maximum likelihood with the ‘mgcv’ R package (Wood 2017). We included only occurrence records collected below 2,500 m of elevation to focus on latitudinal variation on temperature and avoid confounding, regionally-varying effects of elevation (which excluded 3 total occurrence records; Supplemental Table S2).

### Identifying maxima and minima in thermal safety

To identify the latitudes with the highest and lowest thermal safety for each species, we searched for global and local maxima and minima in thermal safety margins, respectively, in the GAM fits. That is, we searched for latitudes in which the first derivative of the fitted function switched from positive to negative, and in which the estimated thermal safety margins were the highest (or lowest) of their neighboring latitudes (*i*.*e*., highest or lowest in a sequence of three). We further confirmed the locations of maximum and minimum thermal safety margins by visually inspecting the plotted GAM fits for each species. To estimate uncertainty, we drew 1,000 samples from the posterior-fitted GAM parameter vector, multiplied by the linear predictors (*i*.*e*., the sum of the product of the predictors and their associated regression coefficients), and detected maxima and minima in each sampled fit, as in Pinsky *et al*. (2019) (Wood 2017; Supplemental Figure S6).

## Results

### Patterns of thermal safety across species and options for avoidance

We estimated that all 8 vector species currently have portions of their distributions in thermal safety and portions in thermal danger (*i*.*e*., positive and negative thermal safety margins, respectively) (Figure 1). Only *Ae. aegypti, An. gambiae*, and *Cx. tarsalis* were thermally safe across the majority of their ranges, with average thermal safety margins of approximately 1°C, (Supplemental Table S4). The remaining species—*Ae. albopictus, An. stephensi, Cx. annulirostris, Cx. pipiens*, and *Cx. quinquefasciatus*—were in thermal danger for most of their distributions with average thermal safety margins ranging from -0.3 to -3.6°C. In particular, *Ae. albopictus, An. stephensi, and Cx. annulirostris* were estimated to be in thermal danger by several degrees Celsius across nearly their entire distributions (Figure 1, Supplemental Table S4).

**Figure 1.**
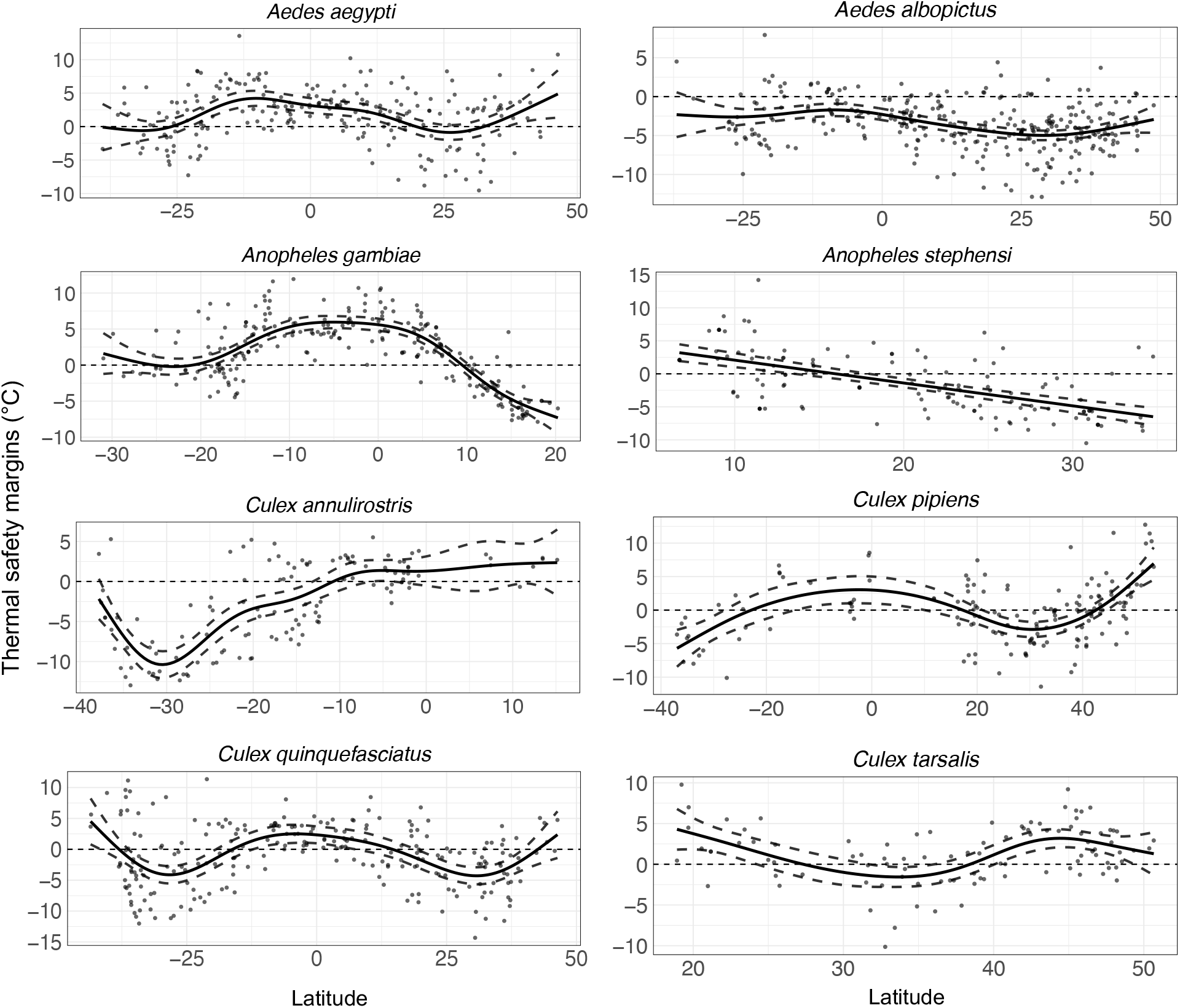
Thermal safety margins across latitude for individual mosquito vectors. Points are estimates at individual occurrence records for a given species; solid and dotted lines show the mean and 95% confidence intervals from the GAM fits. These are estimated with behavioral thermoregulation and with the drought mask, including only observations from below 2,500 m elevation (see Supplemental Figures S3 and S5 for corresponding plots under alternative assumptions).

These results above pertain to estimates under the main model specification, in which options for avoiding thermal extremes—namely, behavioral thermoregulation and reduced activity or aestivation during drought—were included. Thermal safety margin estimates were highly similar when excluding aestivation during drought, but were, on average, 0.13 ± 0.19°C lower under this specification (Supplemental Figure S5, Supplemental Table S4). However, when the ability to behaviorally thermoregulate was removed (*i*.*e*., when vectors were restricted to fully exposed habitats), thermal safety margins were substantially lower (3.15 ± 0.34°C lower) (Supplemental Table S4), and most species were in thermal danger across the majority of their ranges (Supplemental Figure S3), highlighting the importance of access to thermal refugia during high temperature extremes. However, we note that our thermal safety margin estimates use CT_max_ estimated under constant temperatures, and that short-term exposure to higher temperatures may be tolerable, potentially with a physiological cost (as described in the *Methods: Mosquito thermal tolerance data* and Discussion). As expected, thermal safety margin estimates were substantially higher when using CT_max_ estimates from the most thermally tolerant life history trait for a given mosquito species (*e*.*g*., immature survival, biting rate), rather than adult survival for consistency across species (Supplemental Figure S1, Supplemental Table S1). Under this specification, all vector species were thermally safe across the majority of their distributions.

### Latitudinal patterns of vector thermal safety margins

For vector species with distributions spanning both hemispheres, thermal safety margins typically peaked just south of the equator and at the northern temperate range edges (Figure 2, Supplemental Figures S7-S9). For these species, thermal safety margins were lowest between 23 – 35° (N or S)—the latitudinal extent of the subtropics—with the exception of *An. gambiae*, for which thermal safety was lowest in the tropics (∼21°S and 18°N). For the two species with distributions restricted to the northern hemisphere (*An. stephensi, Cx. tarsalis*), thermal safety margins were either highest around the edges of their distribution (∼22°N and 44°N) and lowest around 30-35°N (*Cx. tarsalis*) or peaked at their tropical range edge (∼2°N) and decreased monotonically into the sub-tropics (*An. stephensi*). For all species, these patterns of thermal safety were mostly driven by variation in temperature experienced in fully shaded habitats across latitude (Supplemental Figure S10). That is, the highest hourly mosquito body temperatures were typically estimated for species occurrences in the subtropics, with relatively cooler maximum body temperatures around the equator and northern temperate range edges. These patterns held true when excluding the drought mask (Supplemental Figure S5, Supplemental Table S4).

**Figure 2.**
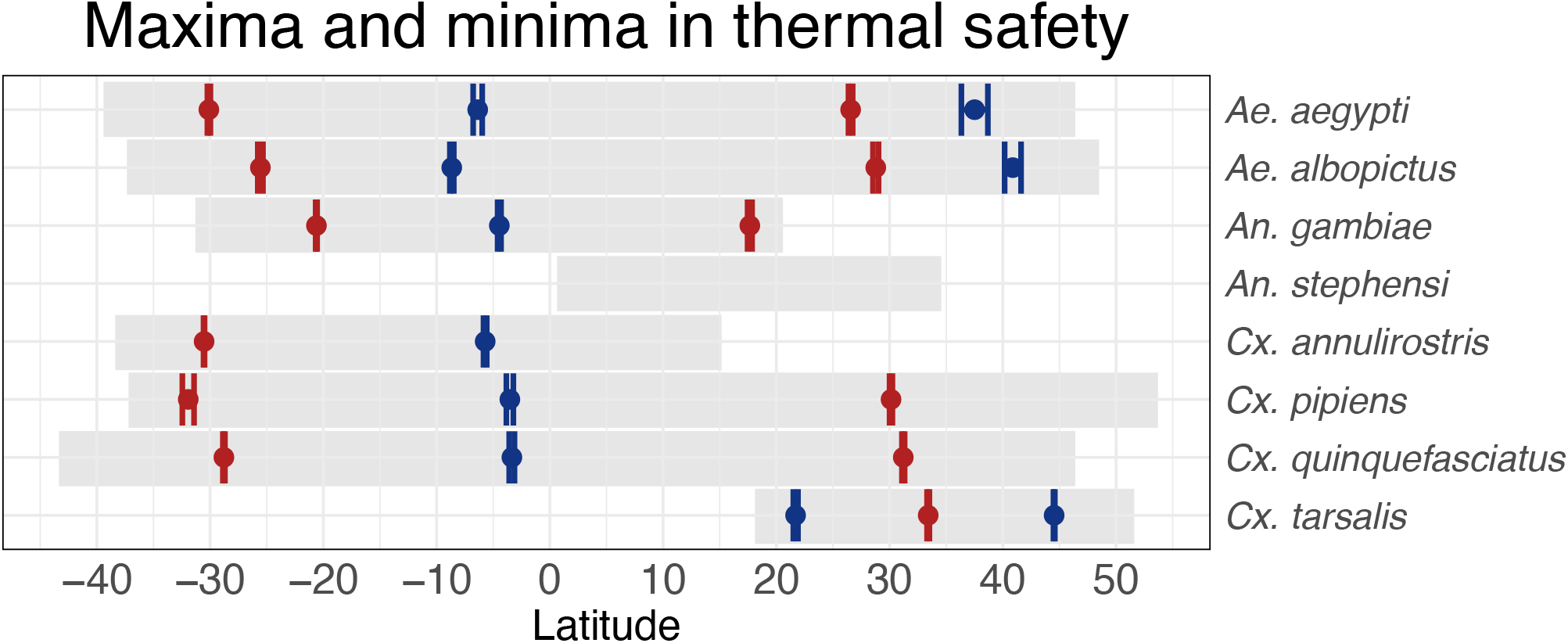
Latitude of maxima (blue points) and minima (red points) in thermal safety margins across the latitudinal ranges (gray rectangles) analyzed for each species. Points and lines denote the mean and 95% confidence interval, respectively, for each species. For *An. stephensi*, no peaks or valleys were identified as the estimated thermal safety margins decrease monotonically across its range. See ‘Methods: *Identifying maxima and minima in thermal safety’* for details on how locations were estimated. See Supplemental Table S3 and Supplemental Figure S6 for estimates and uncertainties.

### Biogeographical patterns of vector thermal safety margins

In addition to thermal safety margins typically being lowest in the subtropics, thermal safety margins varied biogeographically (Figure 3). In particular, regions dominated by deserts and xeric shrublands, including in the Middle East, the western Sahara, and southeastern Australia. had the lowest thermal safety margins across vector species (Figure 3; Supplemental Figure S11) (Olson et al. 2001). Further, the longest consecutive periods of thermal danger (*i*.*e*., when estimated mosquito body temperatures exceed species’ critical thermal maxima; discussed further below) typically occurred in these regions (Supplemental Figure S13).

**Figure 3.**
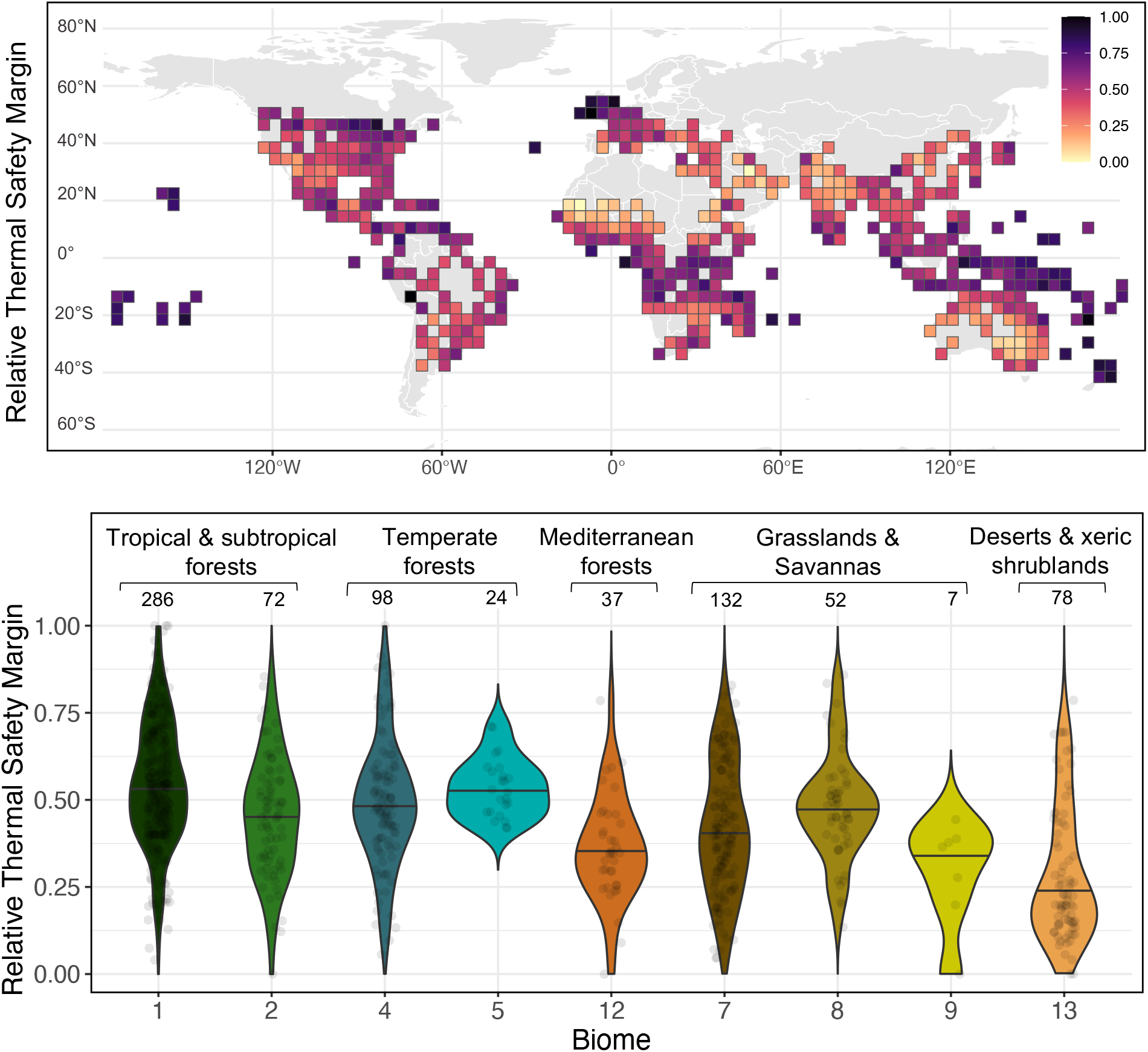
Vector thermal safety margins by grid cell (top) and biome (bottom). For each vector species, thermal safety margins were scaled from 0 to 1 before being combined (by taking the average of each grid cell) as shown here. In the top panel, grid cells are 4°x4°. In the bottom panel, biomes refer to the 14 biomes classified by the World Wildlife Fund. The biome numbers listed on the x-axis correspond to 1) Tropical & Subtropical Moist Broadleaf Forests, 2) Tropical & Subtropical Dry Broadleaf Forests, 4) Temperate Broadleaf & Mixed Forests, 5) Temperate Conifer Forests, 7) Tropical & Subtropical Grasslands, Savannas & Shrublands, 8) Temperate Grasslands, Savannas & Shrublands, 9) Flooded Grasslands & Savannas, 12) Mediterranean Forests, Woodlands & Scrub, 13) Deserts & Xeric Shrublands. Lines within each violin plot denote the median relative TSM for that biome. Points are scaled estimates of TSMs for individual occurrence records within each biome (jittered to improve visibility). The number of occurrence records assigned to each biome is noted above each violin plot. See Supplemental Figures S7-S9 for scaled TSMs for each individual vector species.

### Time scales of thermal danger

The length of time spent in thermal danger (*i*.*e*., when estimated mosquito body temperatures exceed species’ critical thermal maxima), varied widely between species and regions (Figure 4, Supplemental Figure S12, Supplemental Tables S5). In particular, the longest continuous period in thermal danger throughout the year ranged from an average of 1.9 ± 3.4 hours (*Cx. tarsalis*) to 8.0 ± 6.7 hours (*Ae. albopictus*) (mean ± s.d. across occurrence points; Supplemental Figure S12, Supplemental Table S5). Further, the longest period in thermal danger was 8.1 ± 9.4 hours in the subtropics compared to 3.2 ± 4.4 and 4.1 ± 4.5 hours (mean ± s.d. across occurrence records for all species) in tropical and temperate regions, respectively, matching patterns observed in thermal safety margins. Without the ability to behaviorally thermoregulate, the longest period in thermal danger increased by several hours from an average of 4.3 ± 2.1 across all species and regions to 6.1 ± 2.0 (Supplemental Table S5).

**Figure 4.**
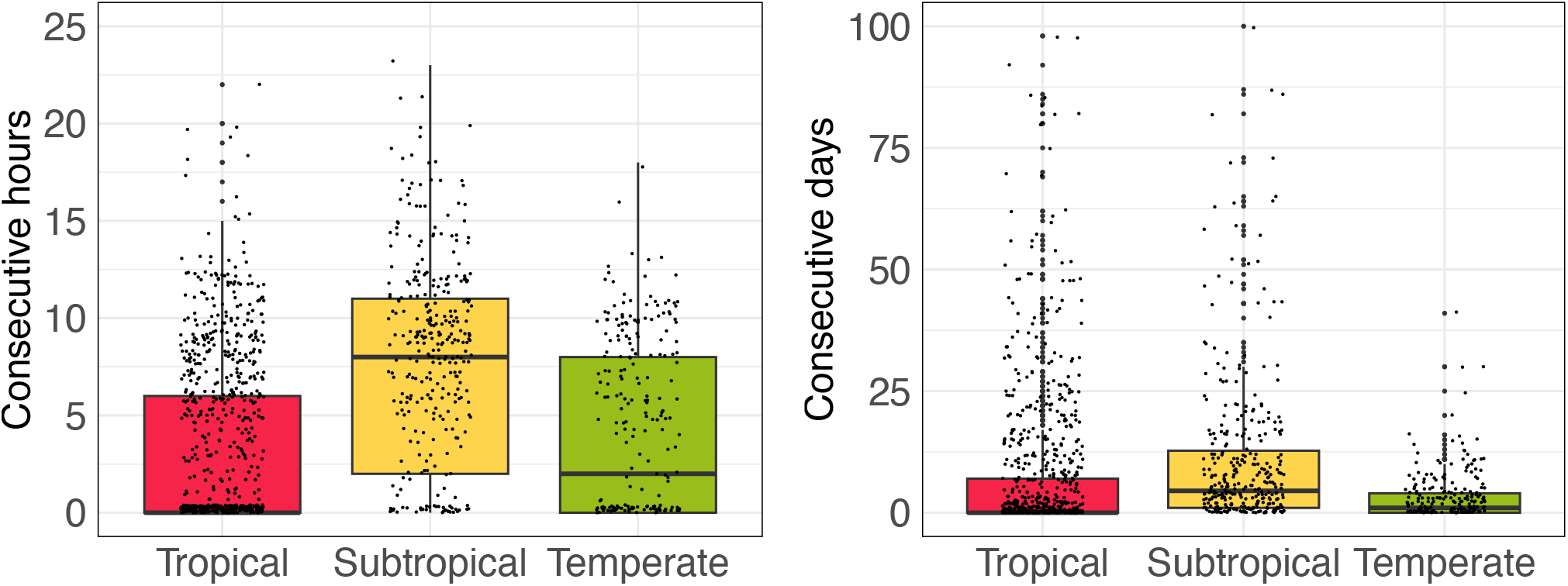
Longest consecutive periods in thermal danger by region for all vector species combined. Y-axes show the longest streak of consecutive hours (left) or days (right) in thermal danger (*i*.*e*., when body temperatures exceed species thermal maxima). The horizontal line within each boxplot denotes the median for that region. See Supplemental Figure S12 and Table S5-S6 for species-specific values of these metrics, and Supplemental Figure S13 for patterns of thermal danger by biome).

These patterns were highly similar when considering the number of consecutive days in which thermal danger occurred for at least one hour. As above, the longest streak of days in thermal danger was lowest for *Cx. tarsalis* (2.1 ± 6.9 days) and highest for *Ae. albopictus* (18.8 ± 27.1 days), and lower in tropical and temperate regions (8.2 ± 18.8, 3.1 ± 5.3 days, respectively) compared to subtropical regions (11.3 ± 17.5 days) (Figure 4, Supplemental Figure S12, Supplemental Table S6). These periods were also substantially longer when the ability to behaviorally thermoregulate was removed, increasing by roughly two-to three-fold for all species. Similarly, when the drought mask was removed, the streaks of thermal danger increased by approximately 2 days (to 9.59 ± 6.07 days) across all species and had a particularly large impact on *An. stephensi* (10.8 ± 15.8 to 17.5 ± 30.6 days) and *Cx. pipiens* (4.1 ± 11.2 to 10.1 ± 30.0 days) (Supplemental Table S6).

## Discussion

We investigated the risk of heat stress posed by climate warming for 8 major mosquito vector species by estimating thermal safety margins—here, the difference between a species’ critical thermal maximum and the hottest hourly body temperature it experiences across its geographic range. We estimated that, when able to access shaded microhabitats, several of the most major vector species, namely *Ae. aegypti, An. gambiae*, and *Cx. tarsalis*, were thermally safe across the majority of their distributions, suggesting low risk from warming (Figure 1). However, when limited to exposed habitats, thermal safety margins for all species were approximately 3°C lower, placing most species in thermal danger (*i*.*e*., body temperature would exceed critical thermal maxima) across large portions of their range (Supplemental Figure S3). Our results thus suggest that behavioral thermoregulation is likely already a pervasive strategy for buffering mosquitoes from high temperature extremes. The reliance on behavior and access to cooler microhabitats to avoid thermal danger has been well documented in other ectotherm species (Kearney *et al*. 2009b; Pinsky *et al*. 2019; Sunday *et al*. 2014). For mosquitoes, behavioral avoidance of high temperatures in laboratory settings has been observed in *Aedes, Anopheles*, and *Culex* species (Blanford *et al*. 2009; Thomson 1938; Verhulst *et al*. 2020), and preference for cooler, shaded oviposition sites in field settings has been documented for *Aedes* (Barrera *et al*. 2006; Vezzani & Albicócco 2009) and *Culex* (Vezzani & Albicócco 2009) species. However, the extent and potential fitness costs of mosquito behavioral thermoregulation remain largely unknown. Our results highlight the importance of understanding mosquito behavioral thermoregulation and of incorporating microhabitat availability when investigating mosquito responses to warming, as behavior could altered projected impacts of future warming (Huey & Kingsolver 2019).

We find that buffering against climate warming for most vector species was highest around the equator and at their northern temperate range edges, and lowest around the subtropics (*i*.*e*., 23-35° N or S) (Figures 1-3). While this finding contradicts the expectation that tropical species are most vulnerable to climate warming from seminal studies such as Deutsch et al. 2008 and Tewksbury et al. 2008, this same pattern was found in a recent meta-analysis including over 400 ectotherm species (Pinsky *et al*. 2019). Similarly, studies of ectotherm range shifts in response to warming, reflecting evidence of climate vulnerability, have found the fastest range shifts at higher latitudes (Ramalho *et al*. 2023), as well as more nuanced responses including east-west and equator-ward shifts (Lenoir & Svenning 2015; Pinsky *et al*. 2013; VanDerWal *et al*. 2013). Our findings thus contribute to a growing body of evidence that species’ risks and responses to climate warming do not vary unidirectionally with latitude, and current risks may be highest in subtropical, rather than tropical, regions because of their higher thermal extremes (Johansson *et al*. 2020; Kingsolver *et al*. 2013; Pinsky *et al*. 2019; Ramalho *et al*. 2023).

Our finding that climate vulnerability is greater in the subtropics is driven by the high short-term temperature extremes experienced there. Across species, we estimated that the highest mosquito body temperatures experienced for one or more hours occurred around 29°N and 38°S (Supplemental Figure S10). This is consistent with expectations from climatology that, although average daily mean temperatures typically peak at the equator and decrease monotonically towards the poles, average daily maximum temperatures typically peak around the subtropics, and are relatively lower at the equator and higher-latitude temperate regions (Buckley & Huey 2016a; Hoffmann 2010; Kingsolver & Buckley 2017; Supplemental Figure S14). These short-term thermal extremes, even if rare, are known to cause major declines in individual fitness, scaling up to drive population and species-level impacts on demographic rates in other ectotherm taxa (Buckley & Huey 2016a, b; Ma *et al*. 2015). Our findings thus suggest that subtropical mosquito populations may be under the greatest pressure to shift their ranges, seasonality, and/or adapt to warming in coming decades, with relatively higher stability in tropical and temperate populations. However, this will depend on the relative fitness costs of exposure to high short-term thermal extremes versus high mean temperatures, which is not well understood and constitutes a key future research direction (Bates & Morley 2020)

While we describe patterns of mosquito climate vulnerability across broad spatial scales (*i*.*e*., between tropical, subtropical, and temperate regions) and latitude, regional-scale biogeographical differences clearly drive variation in vulnerability (Figure 3). We found that thermal safety margins were lowest in specific regions including the Middle East, the western Sahara, and southeastern Australia, which are largely classified as desert and xeric shrubland biomes (Olson *et al*. 2001), but are highly ecologically and climatologically distinct from each other. As described for the subtropics broadly, these regions each experience the highest short-term thermal extremes, although not necessarily the warmest annual mean temperatures (Supplemental Figure S14). Further, while we assumed all locations contained the full range of microhabitat conditions (*i*.*e*., full sun to full shade), the availability of thermal refugia, such as logs, tree holes, and epiphytes, varies widely by region (Scheffers *et al*. 2014), and is likely lower in desert and xeric shrubland biomes, further compounding the higher climate vulnerability there. Humidity, which is important for mosquito survival and may modulate temperature sensitivity, may also limit tolerance of hot temperature extremes, especially in xeric environments (Brown *et al*. 2023).

As climate vulnerability depends not only on the magnitude of temperature extremes, but also on their duration and frequency, we estimated the longest continuous periods that mosquito species spent in thermal danger. As before, we found that the longest stretches of thermal danger, in terms of both consecutive hours and days, occurred in subtropical regions (Figure 4).

Comparing across species, we found that thermal danger, when it occurred, typically lasted no more than 2 to 8 hours (for *Cx. tarsalis* and *Ae. aegypti* to *Ae. albopictus*, respectively, Supplemental Table S5, Supplemental Figure S12). At these time scales, rapid hardening or heat shock responses, which have been well documented in other ectotherm species, could substantially increase short-term critical thermal maxima (Kellermann *et al*. 2017; King & MacRae 2015; Ma *et al*. 2021). As heat shock responses are highly conserved across ectotherm species (Lindquist & Craig 1988), this may be a pervasive strategy for mosquitoes mitigating thermal danger under current and future climatic extremes. However, the precise time scales over which heat shock responses operate, and their overall fitness costs, remain poorly understood. In other ectotherm species, the production of heat shock proteins has been associated with decreased critical thermal maxima on subsequent days (Bai *et al*. 2019) and overall decreases in development and reproduction (Feder & Hofmann 1999; McMillan *et al*. 2005; Sørensen *et al*. 2003). Further, heat injury may be additive, such that recurring exposure to thermal stress may further reduce individual fitness and population growth, particularly if there is insufficient time (*e*.*g*., <6 h) for injury repair (Jørgensen *et al*. 2021; Kashmerry & Bowler 1977). We estimated that certain vector species – namely *Ae. albopictus, Ae. stephensi*, and *Cx. annulirostis –* may experience nearly 10 or more consecutive days of thermal danger in portions of their distribution, which would likely exceed the capacity for heat injury repair and result in reduced fitness (Supplemental Table S6, Supplemental Figure S12). However, other species such as *Ae. aegypti* and *Cx. tarsalis* may experience few consecutive days (*e*.*g*., <3) of thermal stress, potentially enabling recovery and mitigating impacts on fitness. Understanding the impacts of heat stress on mosquitoes at these time scales and frequencies will improve estimates of mosquito climate vulnerability and is an important direction for future research.

The time dimension of heat stress is also important for accurately estimating species critical thermal maxima (Bates & Morley 2020). In this study, we used estimates of critical thermal maxima derived from laboratory experiments, which measured adult mosquito survival under constant temperatures, typically occurring for several hours, days, or weeks (Supplemental Table S1; summarized in Mordecai et al. 2019 and Villena et al. 2022). Prior work has demonstrated that the duration of heat exposure and the rate of temperature change used in heat tolerance assays, as well as the thermal history of the organism itself, including cross-generational effects, can impact the estimated critical thermal maxima (Heerwaarden & Kellermann 2020; Kingsolver *et al*. 2011, 2015; Schiffer *et al*. 2013; Terblanche *et al*. 2007; Waite & Sorte 2022). As a result, there is no single definitive critical thermal maximum for a given taxon or individual, but instead a continuum depending on temporal dynamics of heat exposure and the specific trait of interest (Bates & Morley 2020; Clusella-Trullas *et al*. 2021; Hoffmann *et al*. 2013; Jørgensen *et al*. 2019; Kellermann *et al*. 2012; Lutterschmidt & Hutchison 1997). In general, critical thermal maxima estimated from longer duration heat stress assays, as used here, are lower than those estimated from acute heat exposure assays such as thermal knockdowns (Bates & Morley 2020; Peck *et al*. 2009; Woods *et al*. 2018). Thus, we may have been biased towards calculating smaller thermal safety margins, and thereby overestimating the extent of thermal danger and mosquito climate vulnerability. We used estimates of critical thermal maxima from thermal performance curves derived from constant-temperature laboratory experiments because they are widely available across the focal taxa, they are consistent in their temporal dimension, and they have relevance to fitness in the field (Mordecai *et al*. 2019, 2020; Shocket *et al*. 2018, 2020; Tesla *et al*. 2018). Further, we used the estimate of critical thermal maximum from adult survival for consistency across species, and because this trait is likely the most accurately and precisely estimated, and likely has largest impact on overall fitness.

However, if positive population growth requires all life history traits to be within their thermal limits, we could be underestimating thermal danger. Other measures that could be more locally relevant (*e*.*g*., responses to short-term heat exposures following more realistic daily temperature variation) have not been conducted systematically across taxa and would likely not be broadly representative and comparable across species entire ranges, as we have examined here.

While we sought to provide an ecologically realistic estimate of mosquito risk to climate warming, there are several additional limitations in our risk metric. First, mosquito populations may suffer declines before thermal safety falls to zero, given the typically steep drop-off in thermal performance between a species’ thermal optima and maxima (Mordecai *et al*. 2019), and potential trade-offs in mosquito life history traits. Further, although mosquito thermal tolerance can vary between populations (*e*.*g*., Couper *et al*. 2023; Reisen 1995; Ruybal *et al*. 2016), we applied a single prior estimate of mosquito critical thermal maxima to each species across its range, as population-specific estimates are not yet available. Additionally, seasonality in mosquito life cycles and prolonged periods of dormancy may buffer populations from thermal extremes (Lehmann *et al*. 2010). As mosquito seasonality and dormancy is largely driven by precipitation and moisture conditions (Huestis & Lehmann 2014; Lehmann *et al*. 2010; Shocket *et al*. 2021), we attempted to account for this by applying a drought mask, in which periods following 30 or more days of dry surface soils were excluded from our thermal safety margin estimates (this drought mask had minimal impact on our findings). Incorporating regionally-specific, population-level mosquito responses would enable more accurate estimates of climate vulnerability, but these data are not available for most species and locations. In addition to these limitations, the ultimate impacts of climate warming on mosquito species will depend not only on current thermal vulnerability, but on rates of warming, which are expected to be spatially heterogeneous but generally greatest around the poles (Clem *et al*. 2020), as well as changes in other abiotic and biotic drivers (*e*.*g*., drought, humidity, urbanization, vector control measures; Franklinos *et al*. 2019). For example, urbanization or land clearing could compound the impacts of climate warming by eliminating thermal refugia provided by canopy cover (Alkama & Cescatti 2016). Ongoing work on mosquito population responses to temperature in the face of concurrent global changes will help to further refine estimates of warming-associated risk.

Despite these complexities, our findings are largely in agreement with patterns of ectotherm range shifts in response to warming, suggesting that our metric provides meaningful information on mosquito climate vulnerability. Our results suggest that the subtropics may be most likely to experience shifts in the seasonality and/or intensity of mosquito-borne disease transmission due to climate change in coming decades, potentially necessitating shifts in the types and timing of vector control strategies for effective disease prevention. Further, the tropics—which currently experience the highest mosquito-borne disease burden—could remain relatively favorable for transmission under near-term climate warming, highlighting the need for sustained vector control in this region.

## Data accessibility

All data and code used in this project are publicly available on GitHub: https://github.com/lcouper/VectorThermalSafety/

