## Supplemental Information for "How much warming can mosquito vectors tolerate?"

**Supplemental Methods.** *Estimating thermal safety margins*

**Supplemental Figures (Figures S1-S14)**
**Fig S1.** Thermal safety margins estimated using $CT_{max}$ estimates from the warmest trait
**Fig S2.** Location of individual occurrence records
**Fig S3.** Thermal safety margin estimates with and without behavioral thermoregulation
**Fig S4.** Comparison of soil moisture percent and precipitation
**Fig S5**. Thermal safety margin estimates with and without the drought mask
**Fig S6.** Location of peaks and valleys in thermal safety from GAM fits
**Fig S7**. Thermal safety margin estimates at each occurrence record for *Aedes* species
**Fig S8.** Thermal safety margin estimates at each occurrence record for *Anopheles* species
**Fig S9.** Thermal safety margin estimates at each occurrence record for *Culex* species
**Fig S10.** Maximum experienced temperature in fully shaded microhabitats
**Fig S11.** Global map of terrestrial biomes and all occurrence records used in analysis
**Fig S12.** Length of time in thermal danger by vector species
**Fig S13.** Length of time in thermal danger by biome
**Fig S14.** Global estimates of mean and maximum hourly temperature.

**Supplemental Tables (Tables S1-S6)**
**Table S1.** Species thermal limits across life history traits
**Table S2.** Occurrence record metadata
**Table S3.** Latitude of maxima and minima in thermal safety margins
**Table S4.** Average thermal safety margins for each vector species
**Table S5.** Longest streak of consecutive hours in thermal danger for each species
**Table S6.** Longest streak of consecutive days in thermal danger for each species

**Supplemental References**

**Supplemental Methods**

*Estimating thermal safety margins*

For each species, we estimated thermal safety margin at each occurrence record (*i.e.,* a specific latitude and longitude). To estimate smoothed latitudinal gradients of thermal safety, we used generalized additive models (GAM) fit using restricted maximum likelihood estimation with the 'mgcv' R package (Wood 2017). For each species, s, models were of the form: $\mathrm{TSM_s} \sim \mathrm{s(latitude, k = 8)}$. Herein, k indicates the degree of smoothing, and was chosen to capture the observed non-linearities in the relationship between thermal safety and latitude, while not overfitting, based on visual inspection of the data (Wood 2017).

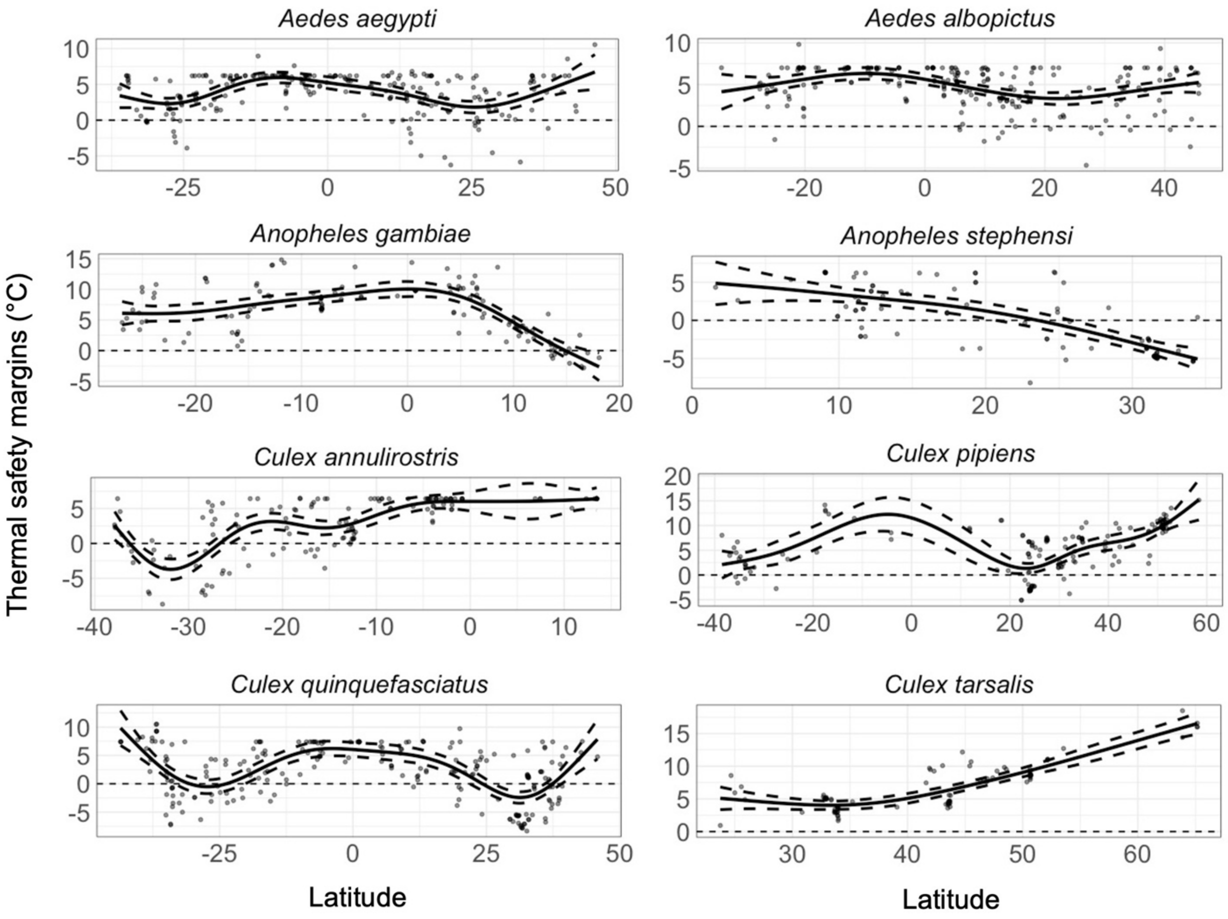

**Supplemental Figure S1.** Thermal safety margins across latitude for individual mosquito vectors estimated using $CT_{max}$ estimates from the most thermally tolerant life history trait (Supplemental Table S1). Points are estimates at individual occurrence records for a given species; solid and dotted lines show the mean and 95% confidence intervals from the GAM fits. These are estimated with behavioral thermoregulation and with the drought mask, including only observations from below 2,500 m elevation.

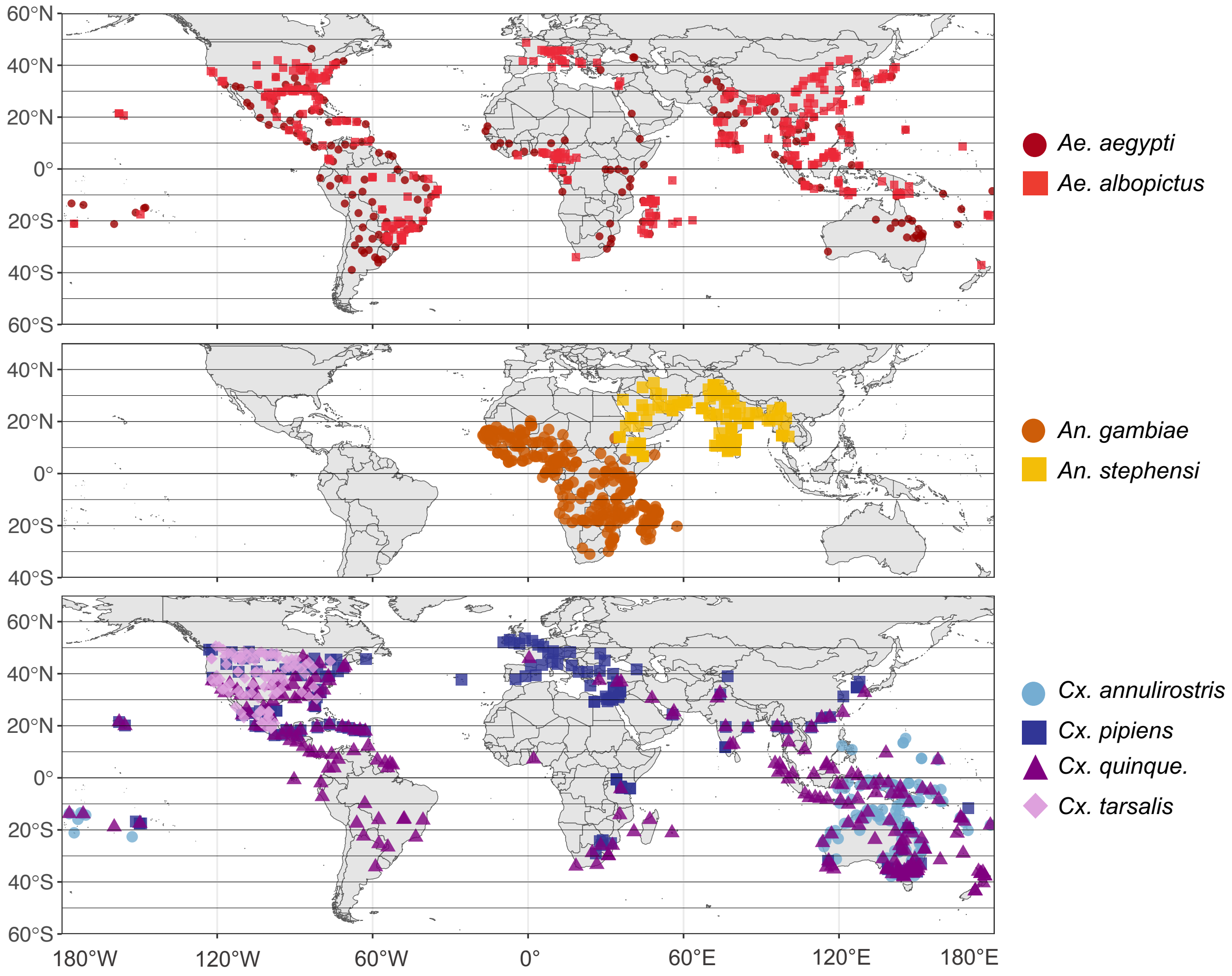

**Supplemental Figure S2.** Location of individual occurrence records for *Aedes* (top), *Anopheles* (middle), and *Culex* (bottom) species used in the analysis. Note that *Cx. quiniquefasciatus* is abbreviated in the legend.

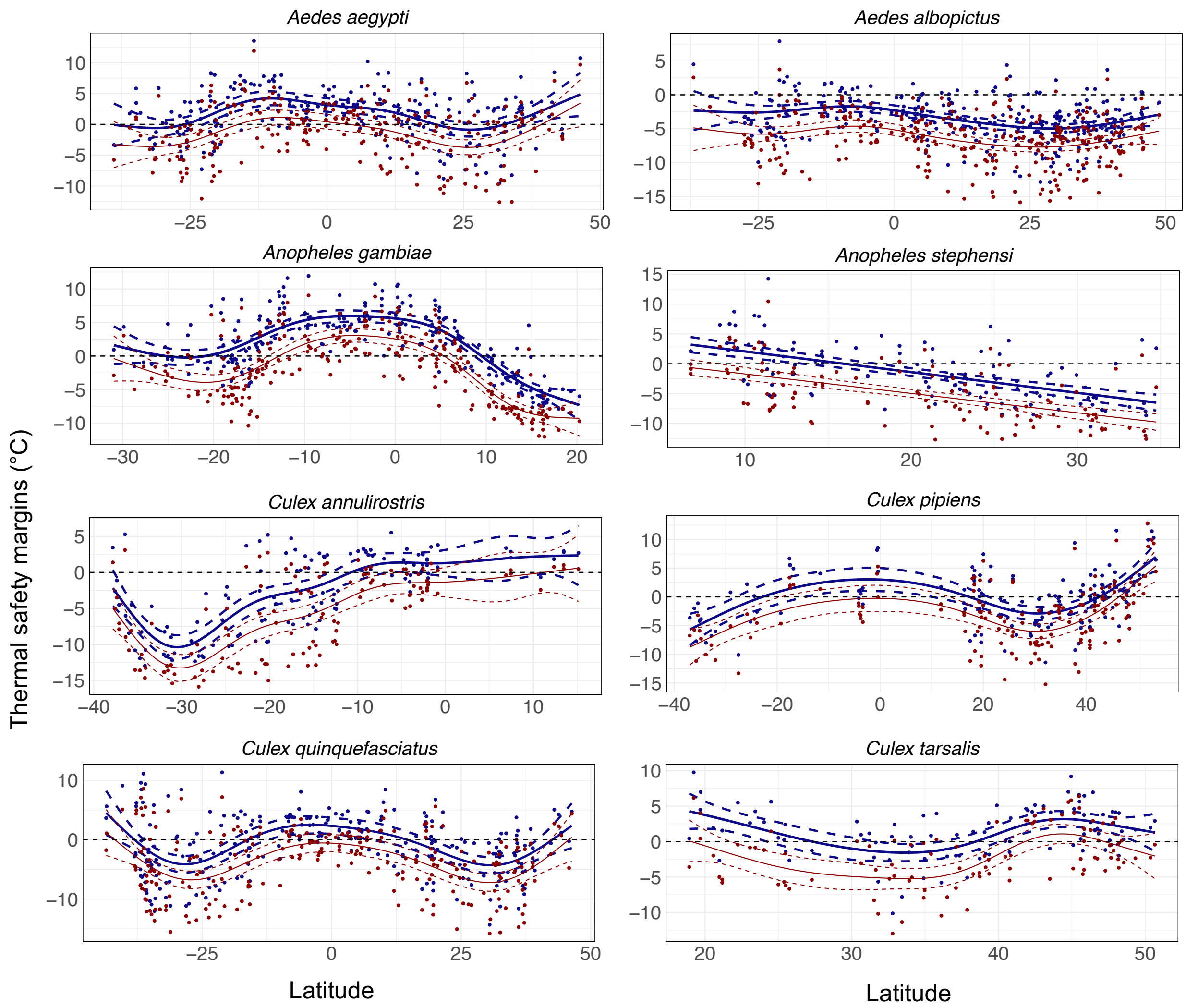

**Supplemental Figure S3.** Thermal safety margins across latitude for individual mosquito vectors estimated with (blue points and lines) and without (red points and lines) behavioral thermoregulation. Points are estimates at individual occurrence records for a given species; solid and dotted lines show the mean and 95% confidence intervals, respectively, from the GAM fits. Note the x- and y-axis limits differ between species.

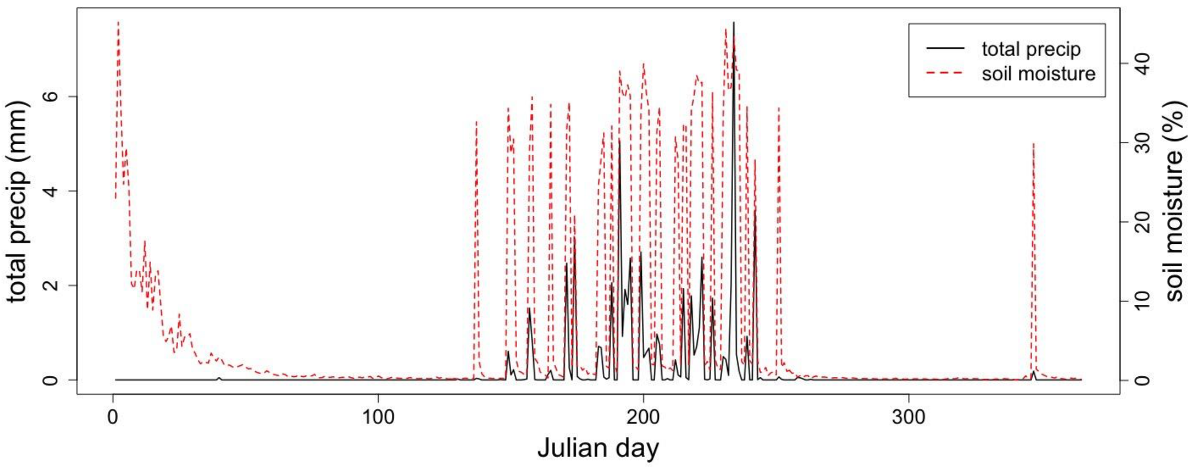

**Supplemental Figure S4.** Comparison of daily soil moisture percent and total precipitation for a given occurrence record (*Anopheles gambiae*; 16.992, 7.979). Soil moisture <5% was used to define the drought mask used in the main analysis.

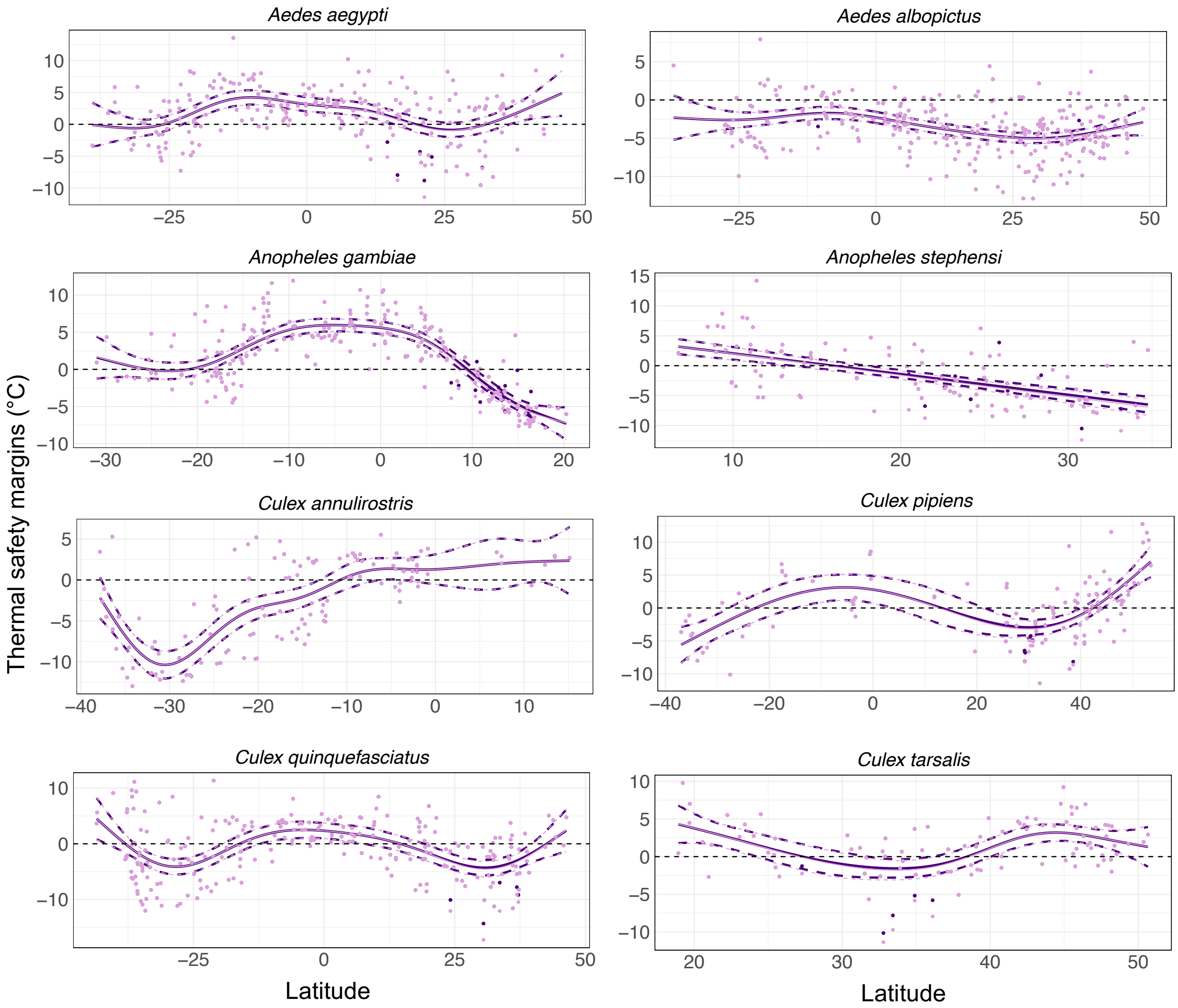

**Supplemental Figure S5.** Thermal safety margins across latitude for individual mosquito vectors estimated with (purple points and lines) and without (pink points and lines) a drought mask (see Methods: *Incorporating impacts of drought on vector life cycles)*. We note that in most cases, the points and lines are nearly entirely overlapping. Points are estimates at individual occurrence records for a given species; solid and dotted lines show the mean and 95% confidence intervals from the GAM fits. Note the x- and y-axis limits differ between species.

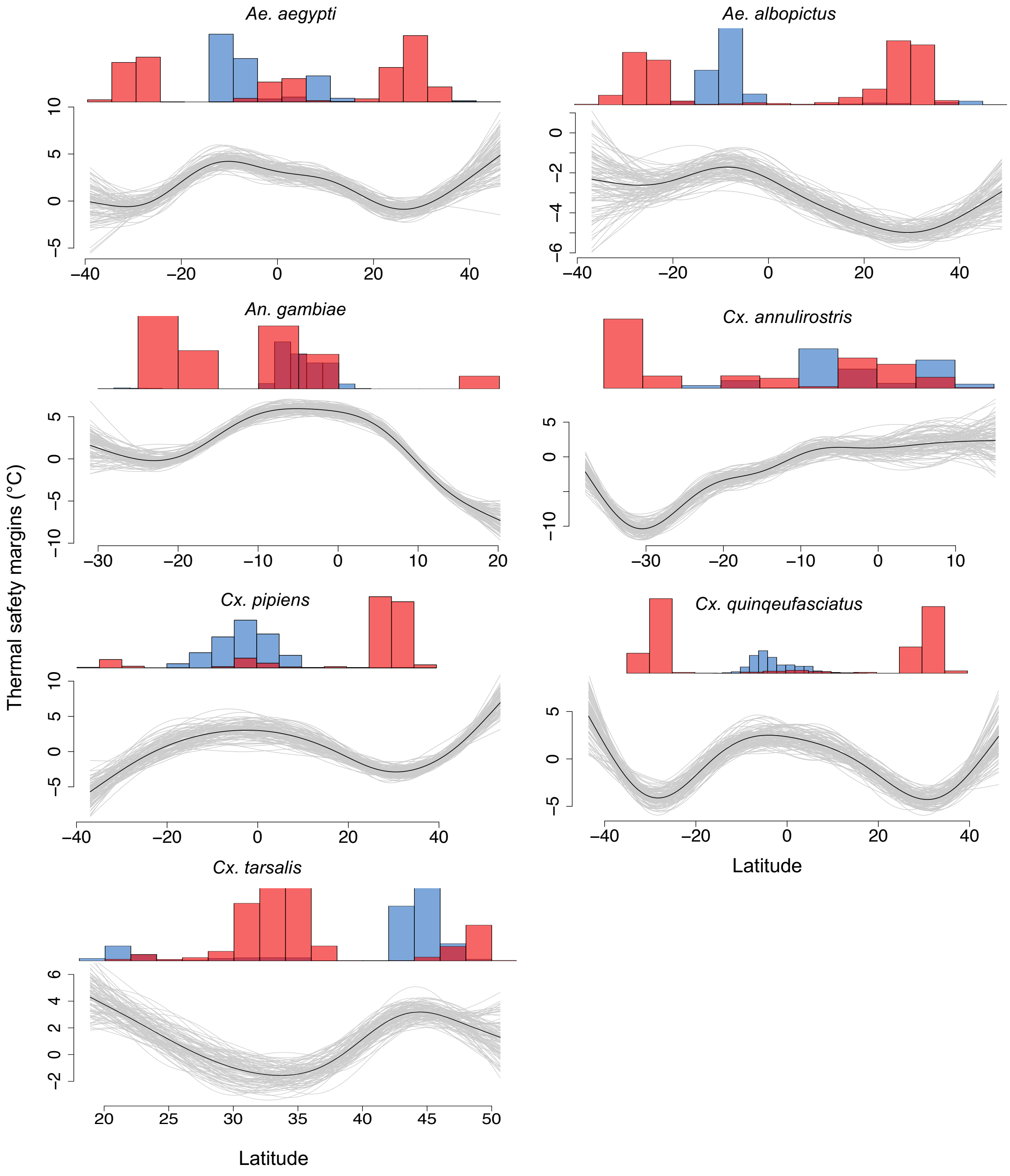

**Supplemental Figure S6.** Uncertainty in the location of peaks and valleys in thermal safety from GAM fits for each species. Line plots for each species show thermal safety across latitude for 100 samples of the posterior-fitted GAM (in gray; mean fit shown in black). Histograms show the spread of estimated peak (blue) and valley (red) locations from the 100 samples. GAM fits are not shown for *An. stephensi* as the estimated thermal safety margins decrease monotonically across its range.

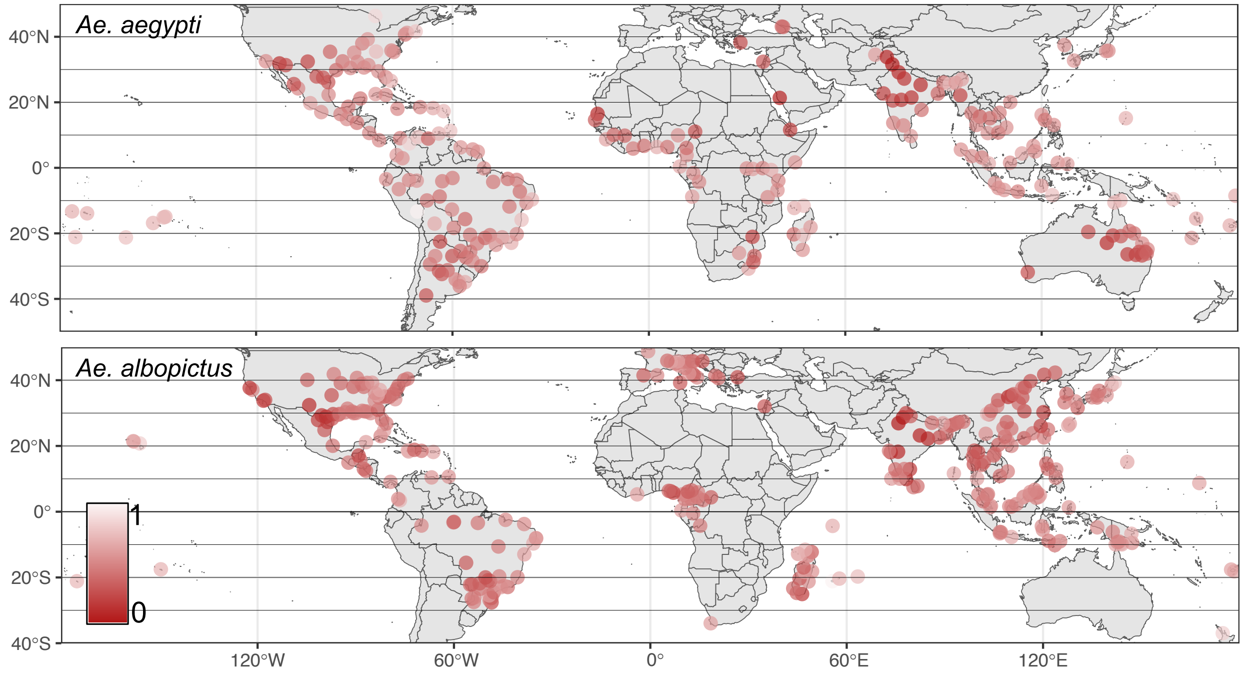

**Supplemental Figure S7.** Estimated thermal safety margins for *Aedes* species at each occurrence record used in the analysis. Here, TSMs are scaled from 0 to 1 for each species (*i.e.*, absolute TSMs are not comparable across species in this plot). The scale bar, which applies for both plots, is shown in the bottom left.

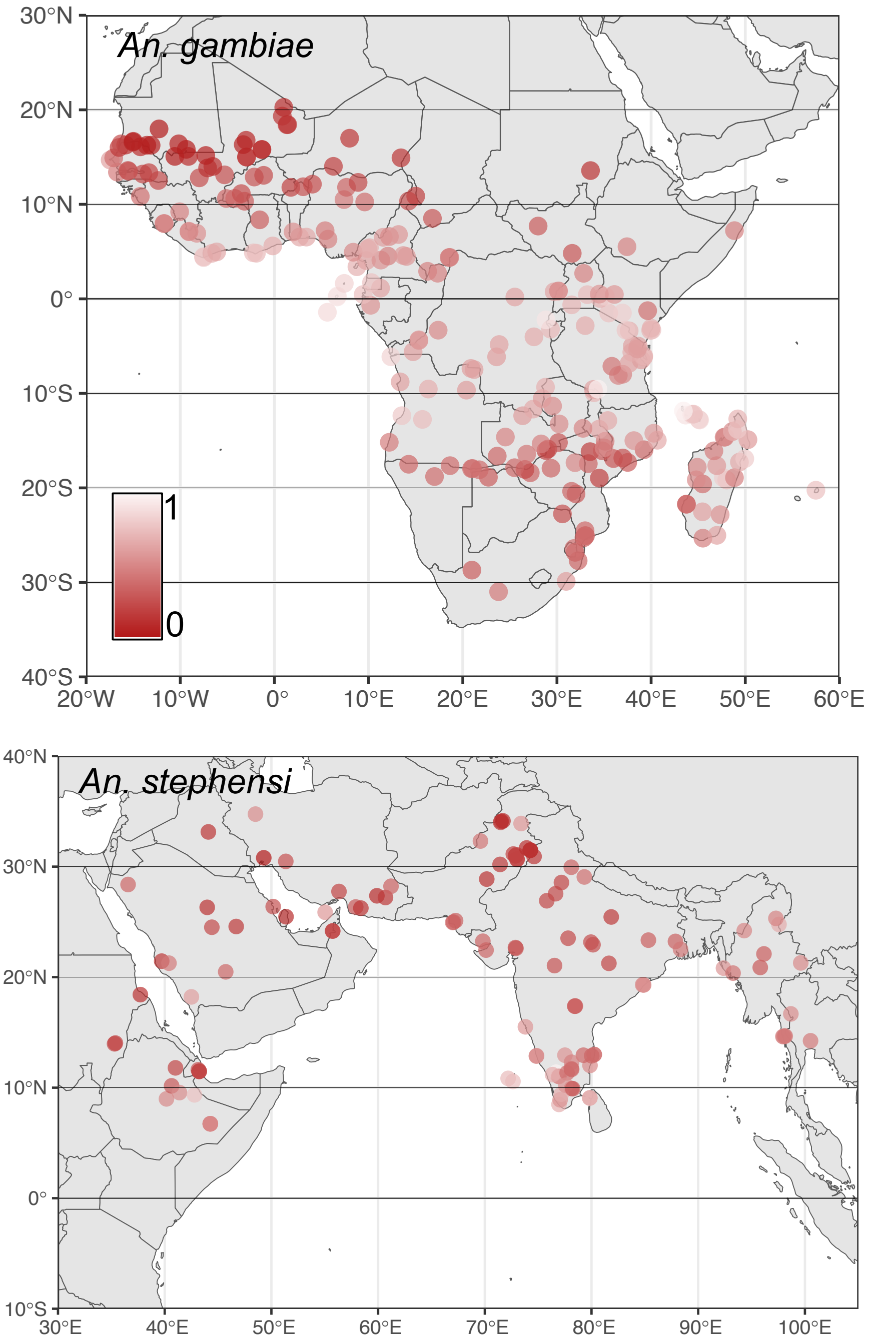

**Supplemental Figure S8.** Estimated thermal safety margins for *Anopheles* species at each occurrence record used in the analysis. Here, TSMs are scaled from 0 to 1 for each species (*i.e.*, absolute TSMs are not comparable across species in this plot). The scale bar, which applies to both plots, is shown in the top plot.

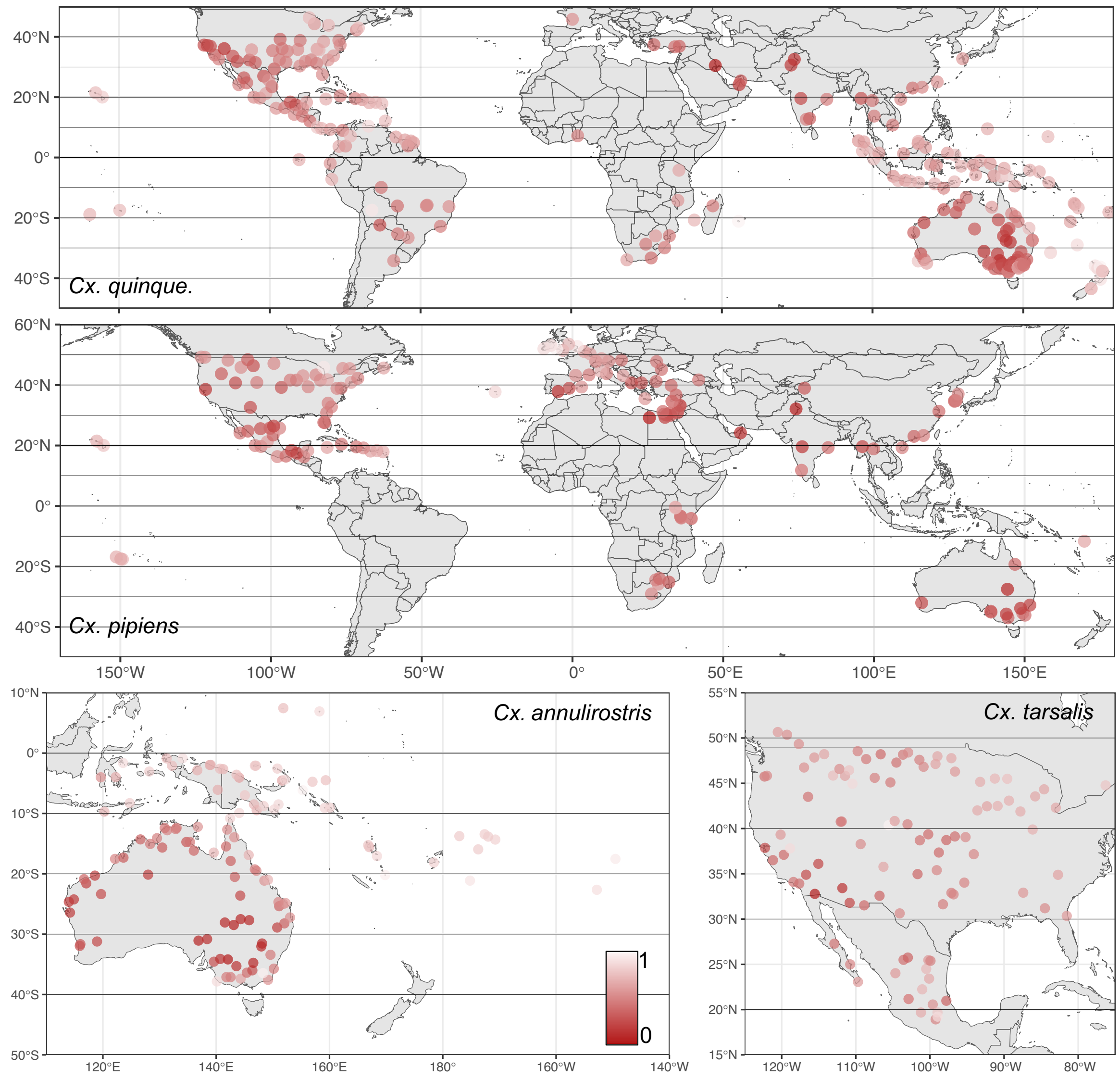

**Supplemental Figure S9.** Estimated thermal safety margins for *Culex* species at each occurrence record used in the analysis. Here, TSMs are scaled from 0 to 1 for each species (*i.e.*, absolute TSMs are not comparable across species in this plot). The scale bar, which applies for all plots, is shown in the bottom center. Note that *Cx. quinquefasciatus* is abbreviated here.

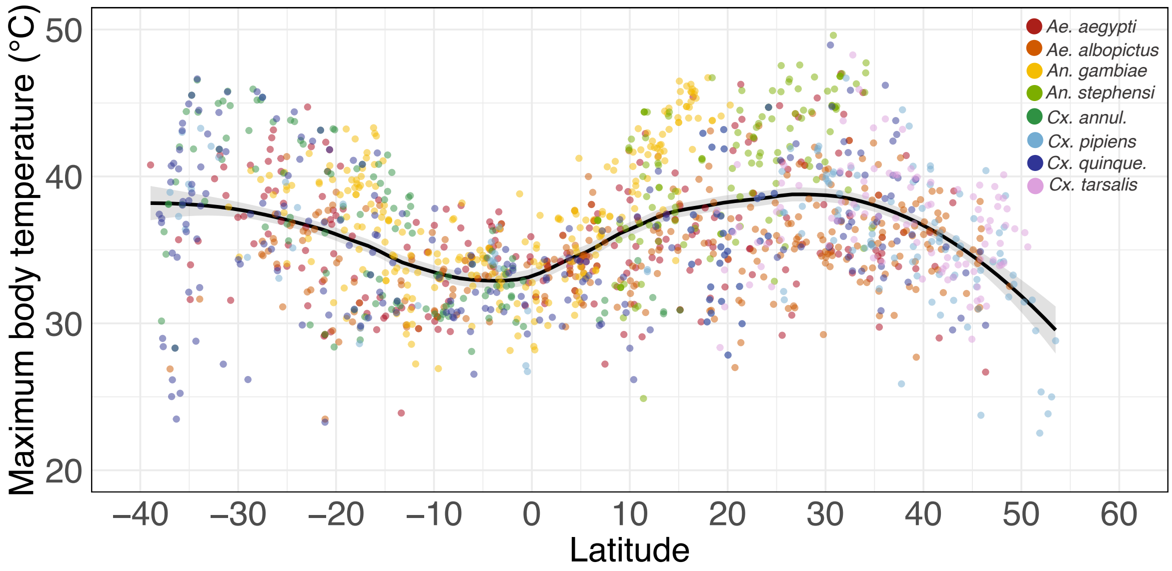

**Supplemental Figure S10.** Maximum experienced temperature in fully shaded microhabitats across latitude. Points are estimates at individual occurrence records and are colored by species; lines and shaded bands show the smoothed means and ±1 standard error considering all species together. Note that *Cx. annulirostris*, and *Cx. quiniquefasciatus* are abbreviated in the legend.

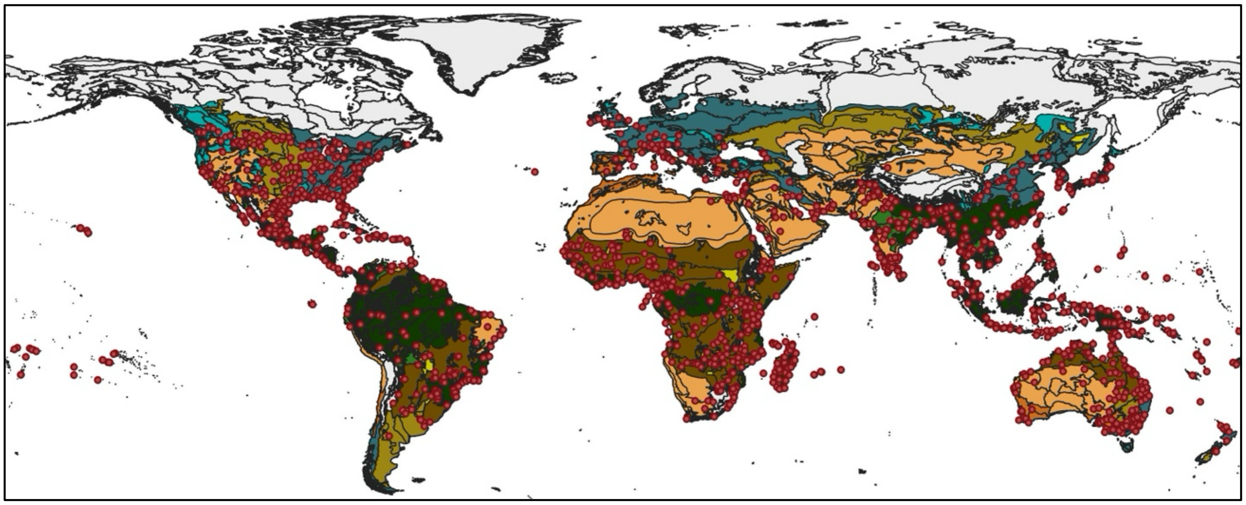

**Supplemental Figure S11.** World map of 14 terrestrial biomes as classified by the World Wildlife Fund (Olson et al. 2001). Occurrence records for any species included in the analysis are shown in light gray circles. The biome numbers listed in the legend correspond to 1) Tropical & Subtropical Moist Broadleaf Forests, 2) Tropical & Subtropical Dry Broadleaf Forests, 4) Temperate Broadleaf & Mixed Forests, 5) Temperate Conifer Forests, 7) Tropical & Subtropical Grasslands, Savannas & Shrublands, 8) Temperate Grasslands, Savannas & Shrublands, 9) Flooded Grasslands & Savannas, 12) Mediterranean Forests, Woodlands & Scrub, 13) Deserts & Xeric Shrublands. 'Other' refers to biomes not used in the analysis as they did not contain any vector occurrence records (*e.g.*, 'Tundra', 'Boreal Forests', 'Mangroves', 'Rock/Ice', 'Lake')

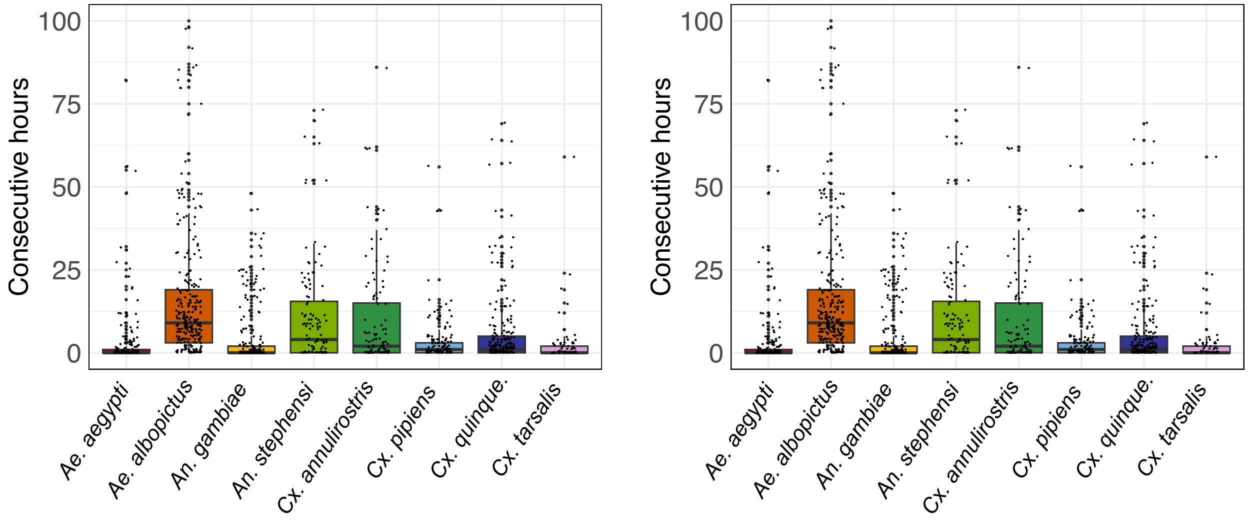

**Supplemental Figure S12.** Longest consecutive periods in thermal danger by vector species. Y-axes show the longest streak of consecutive hours (left) or days (right) in thermal danger (*i.e.*, when body temperatures exceed species thermal maxima).The horizontal line within each boxplot denotes the median for that species.

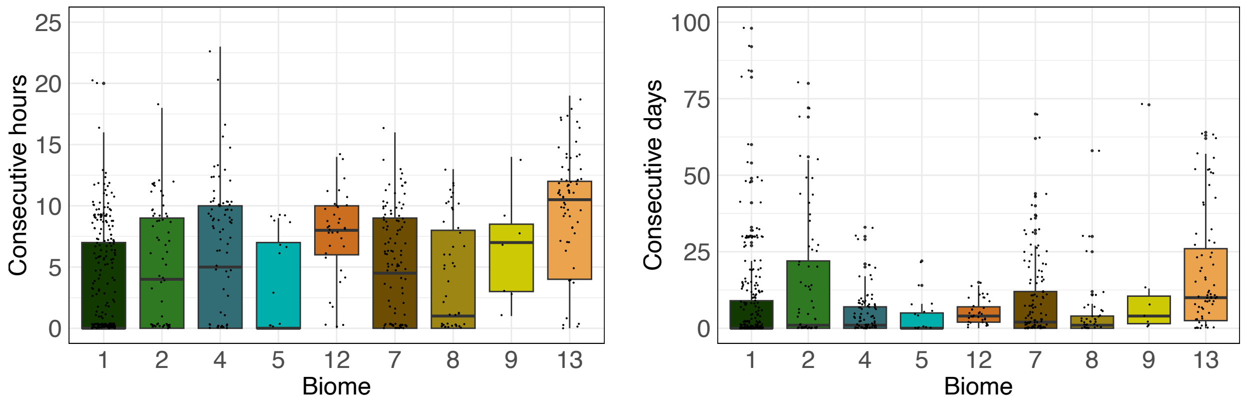

**Supplemental Figure S13.** Longest consecutive periods in thermal danger by biome for all vector species combined. Y-axes show the longest streak of consecutive hours (left) or days (right) in thermal danger (*i.e.*, when body temperatures exceed species thermal maxima). The horizontal line within each boxplot denotes the median for that species. biomes refer to the 14 biomes classified by the World Wildlife Fund. The biome numbers listed on the x-axis correspond to 1) Tropical & Subtropical Moist Broadleaf Forests, 2) Tropical & Subtropical Dry Broadleaf Forests, 4) Temperate Broadleaf & Mixed Forests, 5) Temperate Conifer Forests, 7) Tropical & Subtropical Grasslands, Savannas & Shrublands, 8) Temperate Grasslands, Savannas & Shrublands, 9) Flooded Grasslands & Savannas, 12) Mediterranean Forests, Woodlands & Scrub, 13) Deserts & Xeric Shrublands.

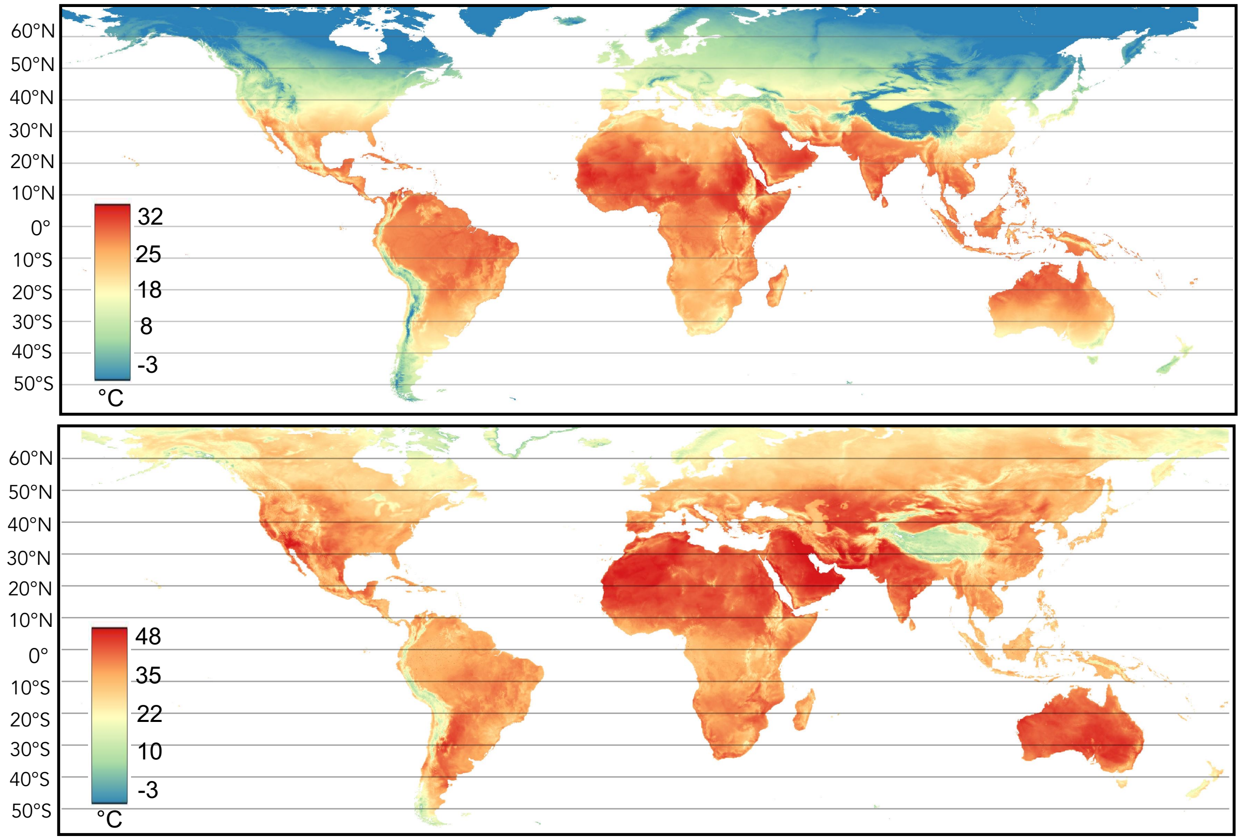

**Supplemental Figure S14**. Global estimates of the mean (top) and maximum (bottom) hourly temperature. Temperature estimates are from ERA5 (the same input as used in the microclimate model). Note that the temperature scales differ between the two plots to better illustrate spatial variation.

**Supplemental Table S1.** Species thermal limits for all available life history traits. Point estimates for critical thermal maximum ($CT_{max}$) and minimum ($CT_{min}$) and thermal optima ($T_{opt}$) are listed for each life history trait as estimated in Mordecai et al. 2019 (for the *Aedes, Culex*, and *Culiseta* species) and Villena et al. 2022 (for the *Anopheles s*pecies). For each species, the most thermally tolerant life history (the history trait with the highest $CT_{max}$, and the trait used in the sensitivity analysis shown in Figure S1) is bolded and denoted with (*). In the main analysis, estimates of thermal performance from adult survival are used. MDR: mosquito immature development rate.

| **Species** | **Trait** | $CT_{min}$ **(°C)** | $T_{opt}$ **(°C)** | $CT_{max}$ **(°C)** |
|---|---|---|---|---|
| *Ae. aegypti* | Adult survival | 10.35 | 23.9 | 37.45 |
| *Ae. aegypti* | Fecundity | 14.7 | 29.6 | 34.4 |
| ***Ae. aegypti*** | **Biting rate*** | **13.8** | **33.8** | **40** |
| *Ae. aegypti* | MDR | 11.6 | 32.7 | 39.1 |
| *Ae. aegypti* | Immature survival | 13.6 | 25.9 | 38.3 |
| *Ae. albopictus* | Adult survival | 13.5 | 22.5 | 31.4 |
| *Ae. albopictus* | Fecundity | 7.9 | 29.4 | 35.6 |
| *Ae. albopictus* | Biting rate | 10.4 | 31.8 | 38.1 |
| ***Ae. albopictus*** | **MDR*** | **8.7** | **32.6** | **39.6** |
| *Ae. albopictus* | Immature survival | 9.1 | 24.2 | 39.3 |
| *Cx. annulirostris* | Adult survival | 13.1 | 23.4 | 33.6 |
| *Cx. annulirostris* | Fecundity | 15.95 | 27.1 | 34.2 |
| *Cx. annulirostris* | Biting rate | 3.8 | 31.8 | 39.1 |
| ***Cx. annulirostris*** | **MDR*** | **11.1** | **32.8** | **39.2** |
| *Cx. annulirostris* | Immature survival | 14.9 | 27 | 39.1 |
| *Cx. annulirostris* | Egg viability | 14.5 | 26.2 | 38 |
| *Cx. pipiens* | Fecundity | 5.3 | 22.1 | 38.9 |
| *Cx. pipiens* | Biting rate | 9.4 | 32.7 | 39.6 |
| *Cx. pipiens* | Proportion ovipositing | 8.2 | 20.8 | 33.2 |
| *Cx. pipiens* | MDR | 0.1 | 30.9 | 38.5 |
| *Cx. pipiens* | Immature survival | 7.8 | 23.1 | 38.4 |
| ***Cx. pipiens*** | **Egg viability*** | **3.2** | **23** | **42.6** |
| *Cx. quinquefasciatus* | Fecundity | 5 | 21.4 | 37.7 |
| ***Cx. quinquefasciatus*** | **Biting rate*** | **3.1** | **31.9** | **39.3** |
| *Cx. quinquefasciatus* | Proportion ovipositing | 1.7 | 24.9 | 31.8 |
| *Cx. quinquefasciatus* | MDR | 0.1 | 31 | 38.6 |
| *Cx. quinquefasciatus* | Immature survival | 8.9 | 23.3 | 37.7 |
| *Cx. quinquefasciatus* | Egg viability | 13.6 | 32.1 | 38 |
| *Cx. tarsalis* | Biting rate | 2.3 | 25.9 | 32 |
| *Cx. tarsalis* | MDR | 4.3 | 32.3 | 39.9 |
| ***Cx. tarsalis*** | **Immature survival*** | **5.9** | **24.6** | **43.1** |
| *An. stephensi* | Adult survival | 6.95 | 21.5 | 39.1 |
| *An. stephensi* | Fecundity | 16.05 | 26.2 | 36 |
| ***An. stephensi*** | **Biting rate*** | **19.1** | **36.05** | **42.25** |
| *An. stephensi* | MDR | 18.6 | 32.2 | 38.05 |

| | | | | |
|---|---|---|---|---|
| *An. stephensi* | Immature survival | 17.7 | 26.1 | 34.7 |
| *An. gambiae* | Adult survival | 8.45 | 22.3 | 38.85 |
| *An. gambiae* | Fecundity | 15.3 | 25.9 | 32.7 |
| ***An. gambiae*** | **Biting rate*** | **34.6** | **16.95** | **43.55** |
| *An. gambiae* | MDR | 13.85 | 29 | 35.95 |
| *An. gambiae* | Immature survival | 16.65 | 24.8 | 32.7 |

**Supplemental Table S2.** Occurrence metadata for each species used in our analysis. This includes the data source for occurrence records, the number of occurrence records analyzed and their latitudinal and elevational ranges. 'GBIF' refers to the Global Biodiversity Information Facility. The number of occurrence records listed below includes only records from <2,500 m (*i.e.,* 1 *Cx. quinquefasciatus* and 2 *Cx. tarsalis* records from higher elevations were removed prior to analysis)

| Species | Occurrence data source(s) | Number records | Latitudinal range (°) | Elevation range (m) |
|---|---|---|---|---|
| *Aedes aegypti* | Kraemer et al. 2015 | 227 | -39.0, 46.4 | -2, 1798 |
| *Ae. albopictus* | Kraemer et al. 2015 | 297 | -37.0, 48.8 | -1, 1784 |
| *Anopheles gambiae* | GBIF.org (23 April 2023) GBIF Occurrence Download https://doi.org/10.15468/dl.7wr3rp Kyalo et al. 2017 | 241 | -31.0, 20.3 | 0, 2022 |
| *An. stephensi* | GBIF.org (23 April 2023) GBIF Occurrence Download https://doi.org/10.15468/dl.ueh9yd Sinka et al. 2020, Ochomo et al. 2023 | 126 | 6.7, 34.8 | -1, 2231 |
| *Culex annulirostris* | GBIF.org (23April 2023) GBIF Occurrence Download https://doi.org/10.15468/dl.m3fces | 123 | -37.8, 15.1 | -6, 2120 |
| *Cx. pipiens* | GBIF.org (23April 2023) GBIF Occurrence Download https://doi.org/10.15468/dl.8qsw69 | 154 | -37.0, 53.5 | -11, 2238 |
| *Cx. quinquefasciatus* | GBIF.org (23April 2023) GBIF Occurrence Download https://doi.org/10.15468/dl.728n59 | 236 | -43.5, 46.3 | -2, 2590 |
| *Cx. tarsalis* | GBIF.org (23April 2023) GBIF Occurrence Download https://doi.org/10.15468/dl.mubyby | 97 | 18.9, 50.7 | -12, 2259 |

**Supplemental Table S3.** Latitude of maxima and minima in thermal safety margins across the latitudinal ranges analyzed for each species. Values denote the estimated location and uncertainty (± 1 se) of maxima and minima in the GAM fits. For *An. stephensi,* no peaks or valleys were identified as the estimated thermal safety margins decrease monotonically across its range. See 'Methods: *Identifying maxima and minima in thermal safety'* for further details.

| Species | Latitudinal Range | Maxima | Minima |
|---|---|---|---|
| *Aedes aegypti* | -39.0°, 46.4° | -6.4 ± 0.4°<br>37.5 ± 1.2° | -30.1 ± 0.1°<br>26.6 ± 0.2° |
| *Ae. albopictus* | -37.0°, 48.8° | -8.7 ± 0.2°<br>40.9 ± 0.7° | -25.6 ± 0.2°<br>28.8 ± 0.3° |
| *An. gambiae* | -31.0°, 20.3° | -4.5 ± 0.2° | -20.6 ± 0.1°<br>17.7 ± 0.2° |
| *An. stephensi* | 6.7°, 34.8° | NA | NA |
| *Culex annulirostris* | -37.8°, 15.1° | -5.7 ± 0.2° | -30.5 ± 0.0° |
| *Cx. pipiens* | -37.0°, 53.5° | -3.5 ± 0.3° | -31.9 ± 0.5°<br>30.1 ± 0.1° |
| *Cx. quinquefasciatus* | -43.5°, 46.3° | -3.4 ± 0.2° | -28.8 ± 0.1°<br>31.2 ± 0.1° |
| *Cx. tarsalis* | 18.9°, 50.7° | 21.7 ± 0.2°<br>44.5 ± 0.1° | 33.4 ± 0.1° |

**Supplemental Table S4.** Average thermal safety margins across the latitudinal range for each vector species under the main model specification (*i.e.*, with behavioral thermoregulation and a drought mask; column 2), without behavioral thermoregulation (column 3), or without the drought mask (column 4). Values denote the mean ± 1 standard deviation.

| **Species** | $\mathbf{TSM_{main\ model}}$ | $\mathbf{TSM_{no\ behavior}}$ | $\mathbf{TSM_{no\ drought\ mask}}$ |
|---|---|---|---|
| *Aedes aegypti* | 1.62 ± 4.06 | -1.34 ± 4.45 | 1.58 ± 4.13 |
| *Ae. albopictus* | -3.61 ± 3.21 | -6.45 ± 3.57 | -3.65 ± 3.24 |
| *Anopheles gambiae* | 1.66 ± 4.59 | -2.04 ± 4.59 | 1.57 ± 4.69 |
| *An. stephensi* | -1.48 ± 4.68 | -5.03 ± 4.70 | -1.68 ± 4.84 |
| *Culex annulirostris* | -2.62 ± 5.18 | -5.81 ± 5.18 | -2.62 ± 5.18 |
| *Cx. pipiens* | -0.34 ± 4.56 | -3.13 ± 4.56 | -0.91 ± 5.01 |
| *Cx. quinquefasciatus* | -1.02 ± 5.01 | -3.93 ± 5.01 | -1.08 ± 5.12 |
| *Cx. tarsalis* | 1.30 ± 3.48 | -1.97 ± 3.48 | 1.26 ± 3.64 |
| Average across species | -0.56 ± 1.99 | -3.72 ± 1.91 | -0.69 ± 1.99 |

**Supplemental Table S5.** Longest streak of consecutive hours in thermal danger by vector species under the main model specification (*i.e.*, with behavioral thermoregulation and a drought mask; column 2), without behavioral thermoregulation (column 3), or without the drought mask (column 4). Values denote the mean ± 1 standard deviation of the longest streak across all occurrence records for a given species (*i.e.*, which may include 0s for records that never experienced thermal danger)

| | $\mathbf{Hours}_{\mathbf{main\ model}}$ | | $\mathbf{Hours}_{\mathbf{no\ behavior}}$ | | $\mathbf{Hours}_{\mathbf{drought\ mask}}$ | |
|---|---|---|---|---|---|---|
| **Species** | Mean | Max | Mean | Max | Mean | Max |
| *Aedes aegypti* | 1.96 ± 3.59 | 14 | 3.42± 3.97 | 15 | 2.01 ± 3.71 | 15 |
| *Ae. albopictus* | 7.99 ± 6.69 | 69 | 9.80 ± 5.89 | 69 | 8.03 ± 6.75 | 69 |
| *Anopheles gambiae* | 2.68 ± 4.11 | 16 | 4.33 ± 4.30 | 16 | 2.81 ± 4.27 | 16 |
| *An. stephensi* | 5.26 ± 4.88 | 14 | 7.03 ± 4.51 | 15 | 5.58 ± 5.18 | 15 |
| *Culex annulirostris* | 5.72 ± 6.06 | 21 | 7.19 ± 5.64 | 21 | 5.72 ± 6.06 | 21 |
| *Cx. pipiens* | 3.88 ± 5.72 | 51 | 5.94 ± 5.62 | 51 | 4.25 ± 4.98 | 51 |
| *Cx. quinquefasciatus* | 4.99 ± 9.06 | 119 | 6.50 ± 8.92 | 119 | 5.22 ± 9.95 | 119 |
| *Cx. tarsalis* | 1.90 ± 3.41 | 16 | 4.59± 4.05 | 17 | 1.90 ± 3.53 | 16 |
| Average across species | 4.30 ± 2.11 | | 6.10 ± 2.02 | | 4.44 ± 2.12 | |

**Supplemental Table S6.** Longest streak of consecutive days in thermal danger (mean ± 1 standard deviation) by vector species under the main model specification (*i.e.*, with behavior and a drought bask; column 2), without behavioral thermoregulation (column 3), or without the drought mask (column 4). Here a day in thermal danger indicates mosquito body temperatures exceeded the species thermal maximum for at least one hour. Values denote the mean ± 1 standard deviation of the longest streak across all occurrence records for a given species (*i.e.*, which may include 0s for records that never experienced thermal danger). Note that *Cx. quinquefasciatus* is abbreviated below.

| | $\mathbf{Days_{main\ model}}$ | | $\mathbf{Days_{no\ behavior}}$ | | $\mathbf{Days_{drought\ mask}}$ | |
|---|---|---|---|---|---|---|
| **Species** | Mean | Max | Mean | Max | Mean | Max |
| *Aedes aegypti* | 2.85 ± 9.32 | 82 | 6.84 ± 15.91 | 141 | 3.42 ± 12.64 | 127 |
| *Ae. albopictus* | 18.76 ± 27.11 | 177 | 31.18 ± 35.84 | 238 | 18.92 ± 27.68 | 177 |
| *Anopheles gambiae* | 4.24 ± 8.90 | 48 | 11.03 ± 18.32 | 80 | 5.24 ± 10.99 | 55 |
| *An. stephensi* | 10.76 ± 15.78 | 73 | 25.06 ± 29.23 | 127 | 17.47 ± 30.55 | 133 |
| *Culex annulirostris* | 9.50 ± 15.93 | 86 | 19.33 ± 29.90 | 145 | 10.94 ± 19.37 | 94 |
| *Cx. pipiens* | 4.06 ± 11.20 | 107 | 8.97 ± 15.14 | 108 | 10.12 ± 29.96 | 167 |
| *Cx. quinque.* | 6.25 ± 13.56 | 108 | 12.11 ± 20.31 | 124 | 7.81 ± 21.87 | 194 |
| *Cx. tarsalis* | 2.07 ± 6.94 | 59 | 5.42 ± 9.49 | 59 | 2.80 ± 10.17 | 81 |
| Average across species | 7.31 ± 5.56 | | 14.99 ± 9.26 | | 9.59 ± 6.07 | |
